# Growth-dependent heterogeneity in the DNA damage response in *Escherichia coli*

**DOI:** 10.1101/2021.05.06.442821

**Authors:** Sebastián Jaramillo-Riveri, James Broughton, Alexander McVey, Teuta Pilizota, Matthew Scott, Meriem El Karoui

## Abstract

In natural environments bacteria are frequently exposed to sub-lethal levels of DNA damage which leads to the induction of a stress response (the SOS response in *Escherichia coli*). Natural environments also vary in nutrient availability, resulting in distinct physiological changes in bacteria which may have direct implications on their capacity to repair their chromosomes. Here, we evaluated the impact of varying the nutrient availability on the expression of the SOS response induced by chronic sub-lethal DNA damage in *E. coli*. The expression of the SOS regulon was found to be highly heterogeneous at the single-cell level in all growth conditions. Surprisingly, we observed a larger fraction of high SOS-induced cells in slow growth as compared with fast growth, despite a higher rate of SOS induction in fast growth. This counter-intuitive result can be explained by the dynamic balance between the rate of SOS induction and the division rates of cells exposed to DNA damage. Taken together, our data illustrates how cell division and physiology come together to produce growth-dependent heterogeneity in the DNA damage response.

## INTRODUCTION

Bacteria are remarkable in their capacity to respond favorably to different environmental conditions, including variations in nutrient availability and perturbations from many different stresses such as oxidative damage or temperature changes. Natural environments vary in their levels of nutrients, affecting the growth of microorganisms. For example, *Escherichia coli* has been estimated to divide every 3 hours inside the intestine, whereas estimates for division time in the urine (bladder) is about 20-30 minutes (Myhrvold *et al*, 2015; Forsyth *et al*, 2018). These variations in growth-rate can have important consequences for how bacteria respond to stresses because they impose constraints on the capacity of bacteria to modify their proteomes in response to changes in the environment (Hui *et al*, 2015). This is particularly true for stresses induced by exposure to antibiotics, as the target of most antibiotics are growth-related processes (Lewis, 2013) and variations in growth-rate correlate with molecular and physiological changes in bacteria (Bremer & Dennis, 2008). For example, the analysis of the interplay between growth-related changes and the response to antibiotics has been useful in gaining quantitative understanding in how bacteria respond to ribosome-targeting antibiotics (Greulich *et al*, 2015). However, how growth-related changes influence the response to other stresses, such as for example DNA damage, has not yet been explored.

DNA damage is one of the most ubiquitous types of stress encountered by bacteria. It can arise from external sources such as exposure to UV light or to DNA damaging agents, for example quinolone antibiotics (Gutierrez *et al*, 2018). Moreover, one major source of DNA damage is directly linked to the cell cycle. Indeed, it has been shown that impaired DNA replication leads to the accumulation of DNA Double Strand Breaks (DSBs) at inactivated replication forks. Spontaneous DSBs have been linked to stalling of the replisome by obstacles, and/or a replication fork encountering DNA nicks and gaps (Michel *et al*, 2018, 2004; Kuzminov, 2001). DNA replication is also involved in the formation of DSBs after exposure to quinolones (Drlica *et al*, 2008; Pohlhaus & Kreuzer, 2005). DSBs are the most deleterious type of DNA damage as they lead to loss of genetic information. They are repaired by homologous recombination where the broken chromosome is repaired using an intact homologous copy as a template. Homology search is catalyzed by RecA which forms a nucleoprotein filament on single strand DNA and promotes strand invasion after a homologous copy has been found. This also leads to the induction of the SOS response (see below).

Changes in growth rate have important consequences on DNA replication in bacteria. In *E. coli*, in rich nutrient conditions, replication of the chromosome is estimated to take about 40 minutes, and segregation/septation to take another 20 minutes, for a nominal cell cycle time of approximately 60 minutes. When cells divide more rapidly than in 60 minutes, they initiate several overlapping rounds of DNA replication (a process referred to as ‘multifork replication’) (Skarstad & Katayama, 2013; Cooper & Helmstetter, 1968). Thus DNA content and the number of replication forks is higher in fast growing cells than in slow growing ones. For example, *E. coli* cells doubling in 30 minutes would contain an average of approximately 12 replication forks-per-cell, while this average drops to approximately 0.36 for cells doubling in 3 hours. Although DSBs are likely to arise more frequently in fast growth conditions because of the high number of replication forks, it is possible that the presence of multiple partial-copies of the chromosome facilitate homology-dependent repair. This raises the question of how DSBs repair may vary with growth conditions. Moreover, growth conditions influence gene expression in bacteria (Hui *et al*, 2015), resulting in less capacity to induce stress-response genes in fast-growth conditions. Thus, bacteria may vary in their capacity to induce the DNA damage response depending on the growth conditions.

*E. coli* responds to DNA damage by inducing expression of the SOS regulon (Radman, 1975; d Ari, 1985; Kreuzer, 2013; Erill *et al*, 2007), which is important for bacteria to survive DNA damaging conditions (Mount *et al*, 1972; Lin & Little, 1988; Mo *et al*, 2016). The SOS regulon is controlled by the LexA transcriptional repressor, which normally binds to SOS promoters thus limiting their transcription. In DNA damaging conditions, LexA binds to the RecA nucleoprotein filaments, resulting in LexA self-cleavage, and leading to the expression of SOS genes (Little, 1991; Butala *et al*, 2011; Kovačič *et al*, 2013). In *E. coli* about 30 genes are under the control of LexA (Fernández De Henestrosa *et al*, 2000), including genes involved in DNA repair (e.g. *recA*), inhibition of cell division (*sulA*), translesion DNA synthesis, toxin-antitoxin modules, and the *lexA* gene itself (Kreuzer, 2013; Baharoglu & Mazel, 2014). In addition to supporting survival under DNA damaging conditions, SOS induction can contribute to increasing mutagenesis (Vaisman *et al*, 2012; Dapa *et al*, 2017; Pribis *et al*, 2019), increasing antibiotic tolerance (Dörr *et al*, 2009; Wu *et al*, 2015), and regulating the transfer of conjugative plasmids and other mobile elements (Beaber *et al*, 2004; Baharoglu *et al*, 2010; Fornelos *et al*, 2016).

Previous reports have shown that SOS expression is heterogeneous in single cells, both in response to DNA damage induced by exogenous agents (Friedman *et al*, 2005; Culyba *et al*, 2018; Uphoff, 2018; Mitosch *et al*, 2019), and when the response is induced by spontaneous DNA damage (Pennington & Rosenberg, 2007; Massoni *et al*, 2012). Heterogeneity in the levels of SOS induction may arise from multiple sources of variability in single cells, including the degree of DNA damage, intrinsic variability in the processes of DNA repair or induction of SOS genes. This raises the question of whether the potential growth-dependence of the formation of DSBs and the subsequent induction of the DNA damage response may also have an impact on the heterogeneity of the SOS response. Given the many consequences of SOS induction to antibiotic tolerance and resistance, it is therefore necessary to address this question at the single cell level.

In this study we address how variation in growth-rate modulated by nutrient quality influences the SOS expression in single cells under conditions of chronic sub-lethal levels of DNA damage. In all conditions, we found a high degree of heterogeneity in SOS levels. We observed that cells with elevated SOS expression were more frequent in slow-growth conditions. However, using a mother machine microfluidics device we established that the rate of SOS induction is higher in fast growth conditions. This apparent contradiction can be explained by the competition between two distinct subpopulations in growing cultures: one population with elevated SOS expression and very long division times, and a second population with moderate SOS expression and normal division times. Because division rates are highly dependent on nutrient conditions, the disparity in division times is much larger in fast growth condition, thus explaining the lower fraction of high SOS cells in rich nutrient. Our observations highlight that the heterogeneity in division times is an important source of single cell variability in the DNA damage response and is likely to play a role in natural environments, where nutrient availability is highly variable.

## RESULTS

### The fraction of cells with spontaneous high levels of SOS induction increases in slow-growth conditions

As a baseline measurement of the DNA damage response in the absence of any artificial source of DNA damage, we characterized the steady-state levels of SOS expression in cells grown in media with different nutrient composition. We quantified SOS induction using a transcriptional reporter based upon the well-characterized SOS promoter *PsulA* driving the expression of *mGFP* and used fluorescence microscopy to measure the fluorescence-per-area (here referred to as “GFP intensity”) in more than 20,000 cells per condition (Supplementary Table 1). This transcriptional fusion was inserted in an ectopic chromosomal locus of a Wild Type *E. coli* strain (WT, MG1655) also carrying an *mKate* marker under the control of a constitutive promoter (*PtetO1*). To ensure that the population is in a balanced state of exponential growth (Schaechter, 2006), cells were grown for at least 12 generations with multiple dilutions before measurements were taken. We chose three growth conditions with population doubling-rates for the WT strain as follows: 0.6±0.01 (SEM) doublings per hour (M9-glycerol, referred to as M9-gly), 1.04±0.04 doublings per hour (M9-glucose, referred to as M9-glu), and 1.61±0.05 doublings per hour (M9-glucose and amino acids, referred to as M9-glu+aa). Importantly, in the fastest growth condition (with doubling every 37 minutes) cells undergo multi-fork replication.

As expected, in the absence of external DNA damage, the vast majority of cells do not show any SOS induction. The main peak of GFP intensity (as measured by *PsulA-mGFP*; Fig. 1A) is almost indistinguishable from the GFP intensity in a strain that is unable to induce SOS (in the ‘SOS-off’ strain, the *lexA3* mutation makes LexA uncleavable (Lin & Little, 1988). In addition, SOS expression for most of the population is close to the level of GFP auto-fluorescence given our imaging conditions (approximately 15-20% difference), consistent with strong repression by LexA acting on the *PsulA* promoter (Supplementary Figure 1).

**Figure 1:**
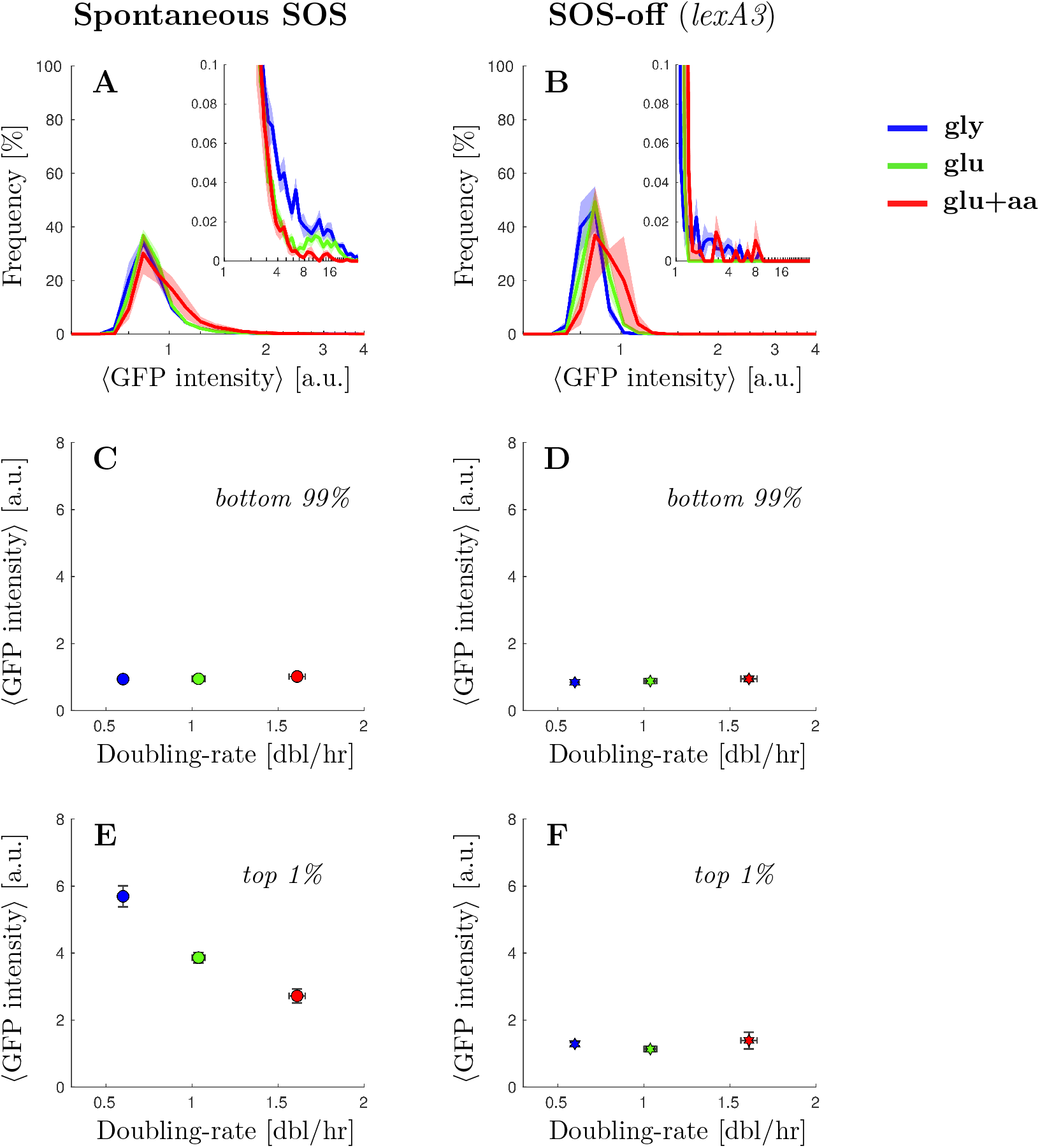
Spontaneous cells with elevated SOS expression are more frequent in low nutrient conditions. A) Steady state distribution of GFP intensity from SOS reporter *PsulA-mGFP* for WT cells in different growth conditions (blue M9-gly, green M9-glu, red M9-glu+aa). GFP fluorescence intensity was measured as arbitrary units of fluorescence per unit cell area. Solid line represents the average frequency and shaded area the standard error from at least 3 replicates done in different days. Inset: A magnification of the second peak at high SOS expression. B) Steady state distribution of GFP intensity from SOS reporter *PsulA-mGFP* for a strain unable to induce SOS (SOS-off, *lexA3* background) in different growth conditions. Solid line represents the average frequency and shaded area the standard error from at least 3 replicates done in different days. Inset: A magnification of the high-fluorescence range of the distribution. C) Average GFP intensity for the lower 99% of the population of WT cells as a function of growth rate. Points represent the average and bars the standard error from biological repeats. D) Average GFP intensity for the lower 99% of the population of *lexA3* cells unable to induce SOS (SOS-off, lexA3 background) as a function of growth rate. Stars represent the average and bars the standard error from at least 3 replicates done in different days. E) Average GFP intensity for the top 1% of the population of wild type cells as a function of growth rate. Points represent the average and bars the standard error from at least 3 replicates done in different days. F) Average GFP intensity for the top 1% of the population of *lexA3* cells unable to induce SOS (SOS-off, *lexA3* background) as a function of growth rate. Stars represent the average and bars the standard error from at least 3 replicates done in different days.

We noticed however that the WT strain has a small subpopulation of highly-expressing cells (Fig. 1A, inset) that is not present in the ‘SOS-off’ mutant (Fig. 1B, inset). Importantly, both the magnitude and location of the secondary peak exhibit strong growth rate dependence. Comparing the blue curve (slow-growth conditions) with the red curve (fast-growth conditions) in Fig. 1A (inset), the fraction of cells in the high SOS state is approximately 6-times higher in slow-growth conditions (cells above 5 A.U. of GFP intensity represent 0.3% and 0.05% for the blue and red curves respectively). We compared the average SOS expression for the highest-expressing 1% of the WT population across growth conditions, and observed that the expression level is higher in the slow-growth condition (5.9±0.4 A.U.) compared to fast-growth condition (2.7±0.2 A.U.) (Figure 1E, and supplementary Figure 2). In contrast, this is not the case for the bottom 99% of the population (Figure 1C).

In other words, under conditions of spontaneous damage, SOS-induced cells are more abundant in slow-growth conditions (compare blue and red curves in Figure 1A, inset), contrary to the expectation that in fast-growth conditions cells may experience more damage due to their higher frequency of DNA replication (multifork replication). Interestingly, in the ‘SOS-off’ strain we found no significant high-expression peak across growth conditions, thus the bimodality in the wildtype data is not due to leakiness from the *PsulA* promoter (Figure 1D and F).

### The fraction of cells showing high levels of SOS expression induced by replication-dependent DSBs increases in slow-growth conditions

Spontaneous DNA-damaging events are rare; to further evaluate the influence of growth-conditions on SOS expression we induced chronic artificial DNA damage. We chose to focus on SOS induction under constant sub-lethal levels of DNA-damage which is commonly occurring in natural conditions (Kuzminov, 1999; Andersson & Hughes, 2014). We used a genetic system that mimics natural replication-dependent breaks based upon the site-specific cleavage of palindromic sequences inserted in the bacterial chromosome (Eykelenboom *et al*, 2008; Cockram *et al*, 2015; Amarh *et al*, 2018). Replication-dependent DSBs at a single locus have been shown to have a minimal effect on the growth rate in rich nutrient conditions, and lead to low levels of SOS induction (Darmon *et al*, 2014). To generate moderate levels of SOS induction, we inserted two palindromes (located at opposite arms of the chromosomes) on the chromosome of an *E. coli* K12 strain containing the *PsulA-mGFP* SOS transcriptional reporter (Materials and Methods).

As with the spontaneous damage in the WT, we observed that SOS levels induced by replication-dependent DSBs were highly heterogeneous in single cells. The SOS levels for the majority of the population showed only a moderate induction, as expected from the occurrence of at most two DSBs occurring only once per cell cycle (Figure 2A). Comparing slow-growth to fast-growth conditions, we observed that the main peak of the distributions shifts slightly to the right in faster-growth conditions indicating higher levels of SOS expression for the bulk of the population (compare blue and red curves in Figure 2A and supplementary Figure 3). Indeed the average value for the main population (bottom 85% of the cells) increases slightly (Figure 2C, 1.7±0.2 A.U. in M9-gly against 2.01±0.08 A.U. in M9-glu+aa). This moderate shift towards higher SOS levels for the main population is consistent with the higher number of replication forks in fast-growth conditions leading to more replication dependent DSBs. We also observed higher SOS values in the strain containing both palindromes than in the strains with a single palindrome consistent with a higher number of DSBs in the double palindrome strain (Supplementary Figure 3A and 3B for each single palindrome strain).

**Figure 2:**
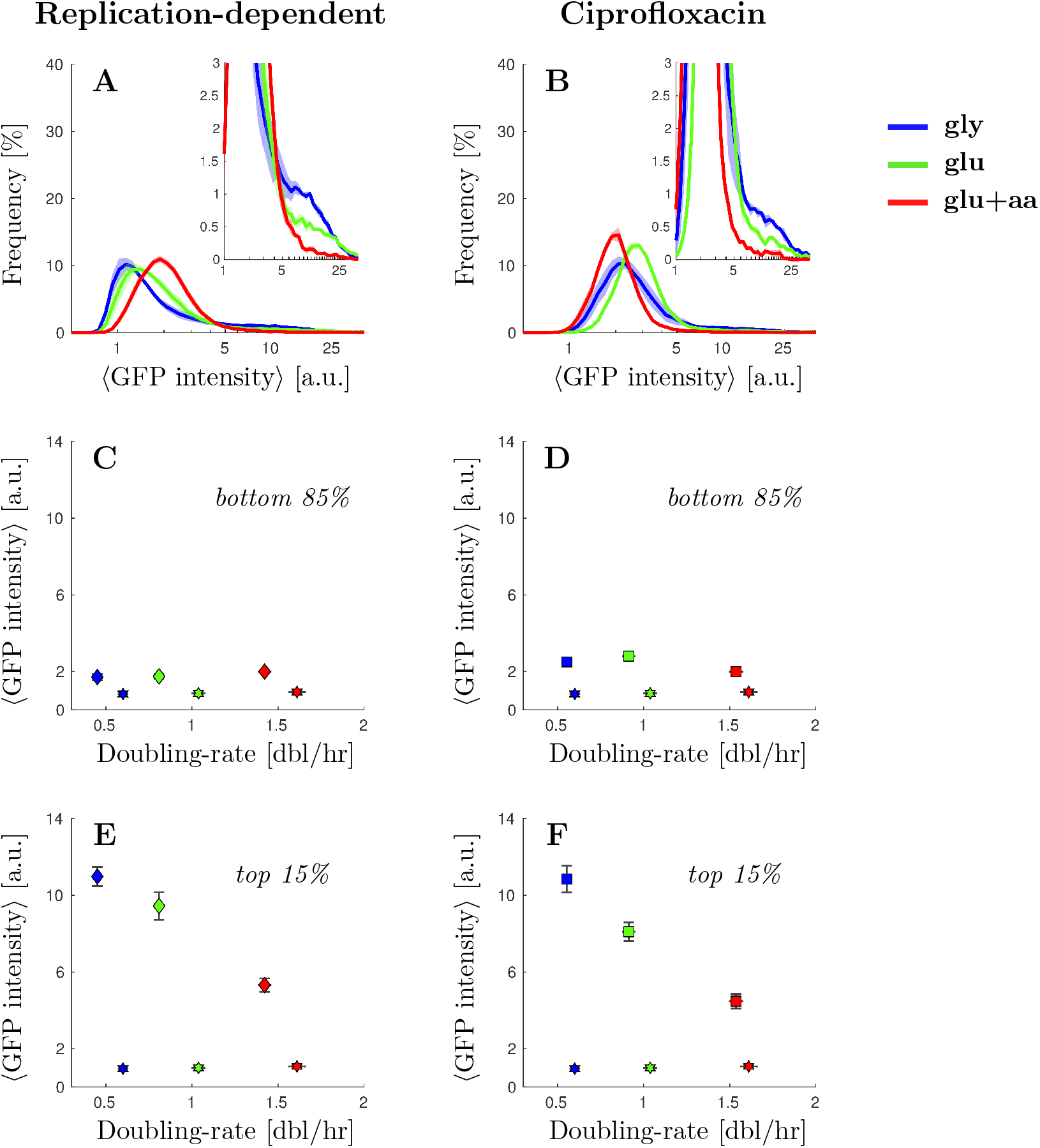
Cells with elevated SOS expression induced by DNA damage are more frequent in low nutrient conditions. A) Steady state distribution of GFP intensity from SOS reporter *PsulA-mGFP* when inducing replication-dependent DSBs (2-pal) in different growth conditions. Solid line represents the average frequency and shaded areas the standard error from at least 3 replicates done in different days. See Supplementary Figures 3A and 3B for the comparable single-palindrome data. Inset: A magnification of the second peak at high SOS expression. B) Steady state distribution of GFP intensity from SOS reporter *PsulA-mGFP* under ciprofloxacin treatment (3 ng/ml) in different growth conditions. Solid line represents the average frequency and shaded area the standard error from at least 3 replicates done in different days. Inset: A magnification of the second peak at high SOS expression. C) Average GFP intensity for the lower 85% of the population under replication-dependent DSBs (2-pal) as a function of growth rate. Diamonds represent the average and bars the standard error from at least 3 replicates done in different days. For comparison, the SOS-off (*lexA3*) mutant data is shown as stars. D) Average GFP intensity for the lower 85% under ciprofloxacin treatment (3 ng/ml) as a function of growth rate. Squares represent the average and bars the standard error from at least 3 replicates done in different days. For comparison, the SOS-off (*lexA3*) mutant data is shown as stars. E) Average GFP intensity for the top 1% of the population under replication-dependent DSBs (2-pal) as a function of growth rate. Points represent the average and bars the standard error from at least 3 replicates done in different days. F) Average GFP intensity for the top 1% under ciprofloxacin treatment (3 ng/ml) as a function of growth rate. Points represent the average and bars the standard error from at least 3 replicates done in different days.

As with spontaneous damage, we observed a secondary peak of high SOS-expressing cells, with a larger fraction of cells in high SOS state in slow-growth condition (Fig. 2A, inset). For example, the fraction of cells whose fluorescence intensity is above 5 A.U. is 14±1% in M9-gly versus 4.3±0.7% in M9-glu+aa. To visualize more clearly this phenomenon, we measured the average SOS levels in the high SOS fractions of the population (Figure 2C and 2E). The average SOS level for the top 15% cells of each population shows a clear negative correlation with population growth rate (Figure 2E, 11.0±0.5 A.U. in M9-gly against 5.3±0.2 A.U. in M9-glu+aa). The SOS levels for the top fraction of the population were also found to be higher in slow-growth conditions for strains carrying single palindromes (Supplementary Figures 3 and 5). This result is unexpected given the positive correlation of SOS levels with growth rates for the main population, suggesting that the high SOS induced population might behave differently than the rest of the population.

### The fraction of cells showing high levels of SOS expression induced by exposure to ciprofloxacin increases in slow-growth conditions

To test the generality of our observation we performed a similar experiment by inducing DSBs using sub-lethal concentrations of a fluoroquinolone, ciprofloxacin (Tamayo *et al*, 2009; Chen *et al*, 1996). As expected, the majority of the population showed a moderate induction in SOS expression which increased with the concentration of ciprofloxacin (1-3 ng/ml; Figure 2B and supplementary Figure 4), The level of induction for the main population did not show any clear growth dependence suggesting that changes in the number of replication forks may not directly relate to the frequency of DNA damage under such low levels of ciprofloxacin exposure.

However, we observed, as previously, growth-dependent heterogeneity in the response. Indeed, a fraction of the population reached very high SOS induction and this fraction was larger in poor than in rich nutrient conditions (Figure 2B, inset). For example 13±2% of the cells show a fluorescence intensity above 5 A.U. in M9-gly versus 2.4±0.3% in M9-glu+aa. Furthermore, under exposure to 3 ng/ml of ciprofloxacin, we observed that the average SOS level for the top 15% cells of each population shows again a clear negative correlation with population growth rate (10.9±0.7 A.U. against 4.5±0.4 A.U. comparing the red and blue squares). In contrast, the bottom 85% shows negligible growth rate correlation (2.5±0.2 A.U. against 2.00±0.06 A.U. comparing the red and blue squares). Similar trends were observed for intermediate doses of ciprofloxacin (Supplementary Figures 4 and 5). Therefore we conclude that, consistent with the other mechanisms of DNA damage studied, exposure to ciprofloxacin leads to a subpopulation of cells with high SOS induction that behave differently from the rest of the population with respect to growth-rate change.

### Cells with high levels of SOS induction arrest division

The data presented so far indicate that, independently of the type of DSBs induction, the fraction of cells with high SOS expression in a snapshot of the population distribution at equilibrium is higher in slow growth conditions. To better understand the dynamic interplay between growth conditions and SOS induction by DNA-damage, we used time-lapse microscopy to observe single-cells growing on agar pads for about eight divisions. As expected, we observed only a small fraction of cells inducing very high levels of SOS expression (Supplementary Figures 6 and 7). The majority of cells with very high SOS levels delayed or stopped division, which is consistent with the induction of the SOS dependent cell division inhibitor SulA (Huisman *et al*, 1984; Cambridge *et al*, 2014; Burby & Simmons, 2019). In contrast, cells with more moderate levels of SOS induction went through several rounds of division during the course of the experiment.

Thus our time-lapse data suggests that there are two subpopulations present: one that divides at a normal rate with relatively low or intermediate SOS expression levels, and a second subpopulation that divides very slowly with high SOS expression levels. This has direct consequences on the relative abundance of each type of cell in a growing population: high SOS cells might be partially out-competed by the rest of the population which is dividing faster than them.

### The transition rate to high-SOS state is higher in fast-growth conditions

To better understand the dynamics of the induction of low and high SOS levels in cells independently of the competition that arises from the differences in division rates in these two states, we made use of a microfluidic mother machine (Wang *et al*, 2010). In this set-up, each cell is trapped in its individual channel where it is possible to measure in real time individual rates of SOS induction and division. We collected fluorescence images of the strain carrying the 2 palindromes over 10 to 40 hours in the three growth media used previously. We used the constitutively expressed *mKate2* marker for cell segmentation and detection of division and the *PsulA-mGFP* marker to monitor SOS induction. Consistent with our observation on agar pads, we observed cell lineages which induced moderate levels of SOS and continued to divide (Figure 3A) as well as lineages where SOS induction was higher and cell division was strongly delayed or arrested leading to the formation of filaments (Figure 3B). These experiments were performed at least three times in each growth condition. The distributions of division time, cell elongation rate and fluorescence intensity are shown in Supplementary Figure 8 and show good day-to-day reproducibility.

**Figure 3:**
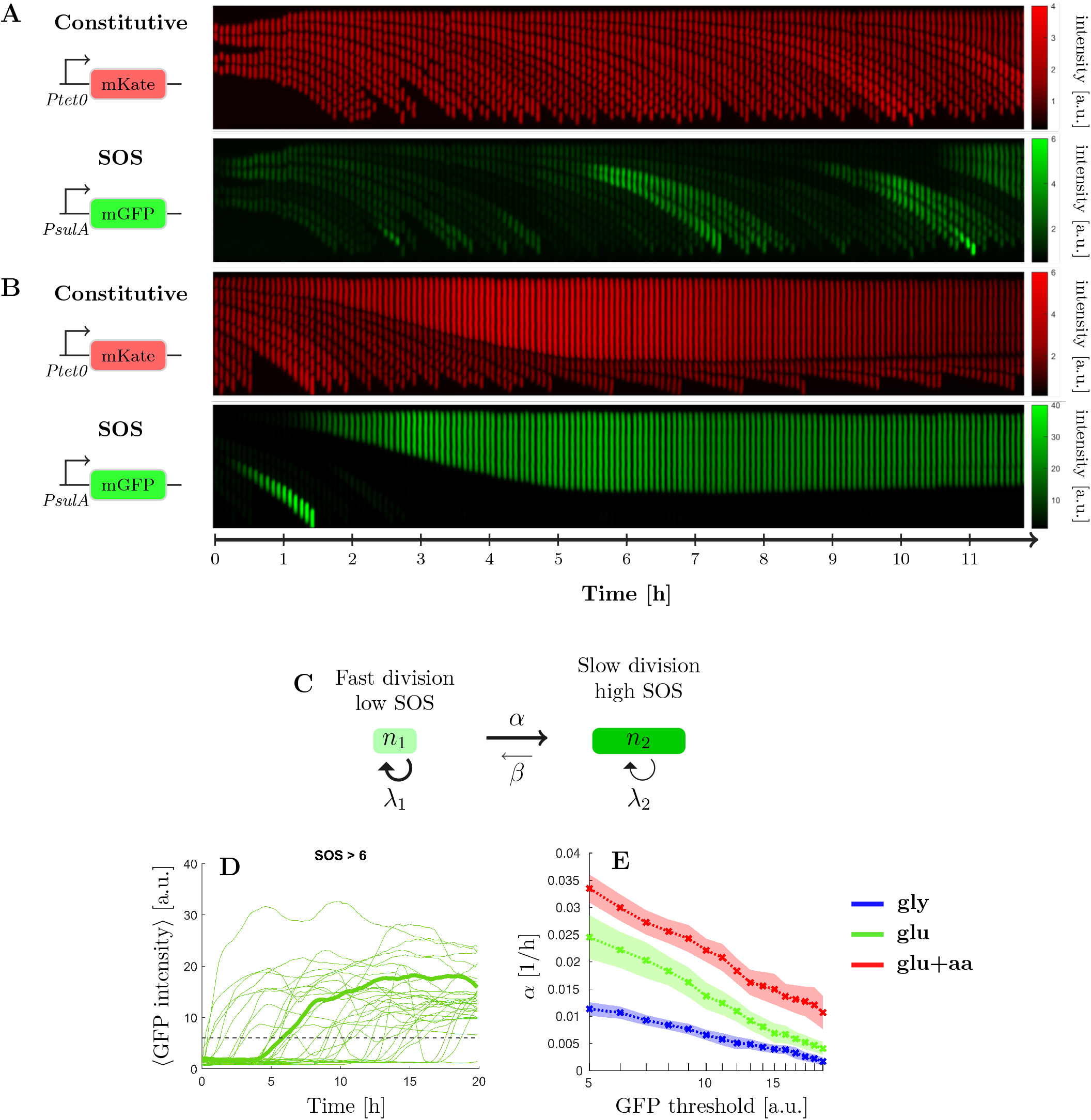
The transition rate to high-SOS state is higher in fast-growth conditions. Representative kymographs of a strain experiencing replication-dependent DSBs (2-pal) in M9-glucose+amino-acids medium. Scale bars represent normalised intensity for constitutive reporter *PtetO-mKate2* and GFP intensity from SOS reporter *PsulA-mGFP*. A) Example mother cell lineage with low level of SOS induction. Top, constitutive expression of *mKate*, bottom *PsulA-mGFP*. B) Example mother cell lineage inducing a high level of SOS. Top, constitutive expression of *mKate*, bottom *PsulA-mGFP*. Cell division is inhibited while cell growth continues and eventually stops. C) Switching model. Cells with low levels of SOS induction rate switch at rate to a high SOS level state and switch back at rate (with *β* ≪ *α*). D) GFP intensity trajectories of SOS induction observed in the 2 palindrome strain as a function of time. One trajectory has been highlighted in bold. The dashed line correspond to the threshold discriminating between high and low-SOS cells. E) Switching rate estimated for multiple GFP intensity thresholds under replication-dependent DSBs (2-pal) in different growth conditions (see supplementary method for the estimation). The switching rate is always higher in rich than in poor nutrient. Points represent the average and the shaded area the standard error from three biological repeats.

To account for the dynamic equilibrium of the high and low SOS populations in steady-state, we adapted a previous mathematical model used to explain the population dynamics of persister cells (Balaban *et al*, 2004; Patra & Klumpp, 2013). In this model, cells can be in either of two states (high and low SOS, Figure 3C), corresponding to different division rates (*λ*_1_ and *λ*_2_). The total number of cells in the population is fixed, as we follow only mother cells in the microfluidics device, and cells can switch from low to high SOS at rate α (switching back at rate β). An example of time traces of cell trajectories for expression of the *PsulA-mGFP* is shown in Figure 3D: we recorded the time at which each cell reached a threshold of GFP intensity above 5 A.U. to estimate the rate at which cells induce high level of SOS. Very few cells reverted from high to low SOS, so we consider β to be negligible. We used Maximum Likelihood estimation (see supplementary methods) to compute α for GFP thresholds ranging from 5 to 20 A.U. in the three growth media (Supplementary Figure 9). As seen in Figure 3E, the rate of switching to high SOS is always higher in rich nutrient condition than in poorer ones irrespective of the threshold, although 5 A.U. is the threshold that gives the highest discriminatory power. This indicates that individual cells have a higher probability per unit of time of switching to high SOS in rich nutrient condition, in keeping with the higher number of replication forks in these conditions. Therefore, the larger fraction of high SOS cells that we observed in slow-growing populations at steady state is *not* explained by a higher rate of SOS induction. Rather, it is the result of a competition between high SOS/slow dividing and low SOS/fast dividing cells.

### A mathematical model based on two competing subpopulations explains the large fraction of high SOS cells observed in slow growth conditions

To better understand the interplay between the rate of SOS induction and the division rates of low and high SOS cells, we expanded our previous model to describe a growing population (Figure 4A). Given that the rate of switching from high SOS to low SOS (β) is very low and that the division rate of high SOS cell (*λ*_2_ is also negligible, we can show that the expected fraction of high SOS cell (*f*_2_) is approximated by 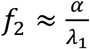 (Figure 4B and supplementary methods for the exact result). We estimated *λ*_1_ from the division rates of the low SOS cells in the mother machine experiments using previously established methods (Painter & Marr, 1968; Thomas, 2017) (see supplementary Figure 8 and supplementary method). We observed that the population growth rate in the mother machine was similar to the batch experiment in M9-glu-aa for the WT strain but lower in M9-gly and M9-glu by approximately 20% (Figure 4C) possibly due to slight constriction of the cells in the mother machine device (Yang *et al*, 2018). This trend was more pronounced in the strain carrying two palindromes which may be due to over-estimation of the batch growth rate due to the impact of filamentation on OD measurements (Stevenson *et al*, 2016).

**Figure 4:**
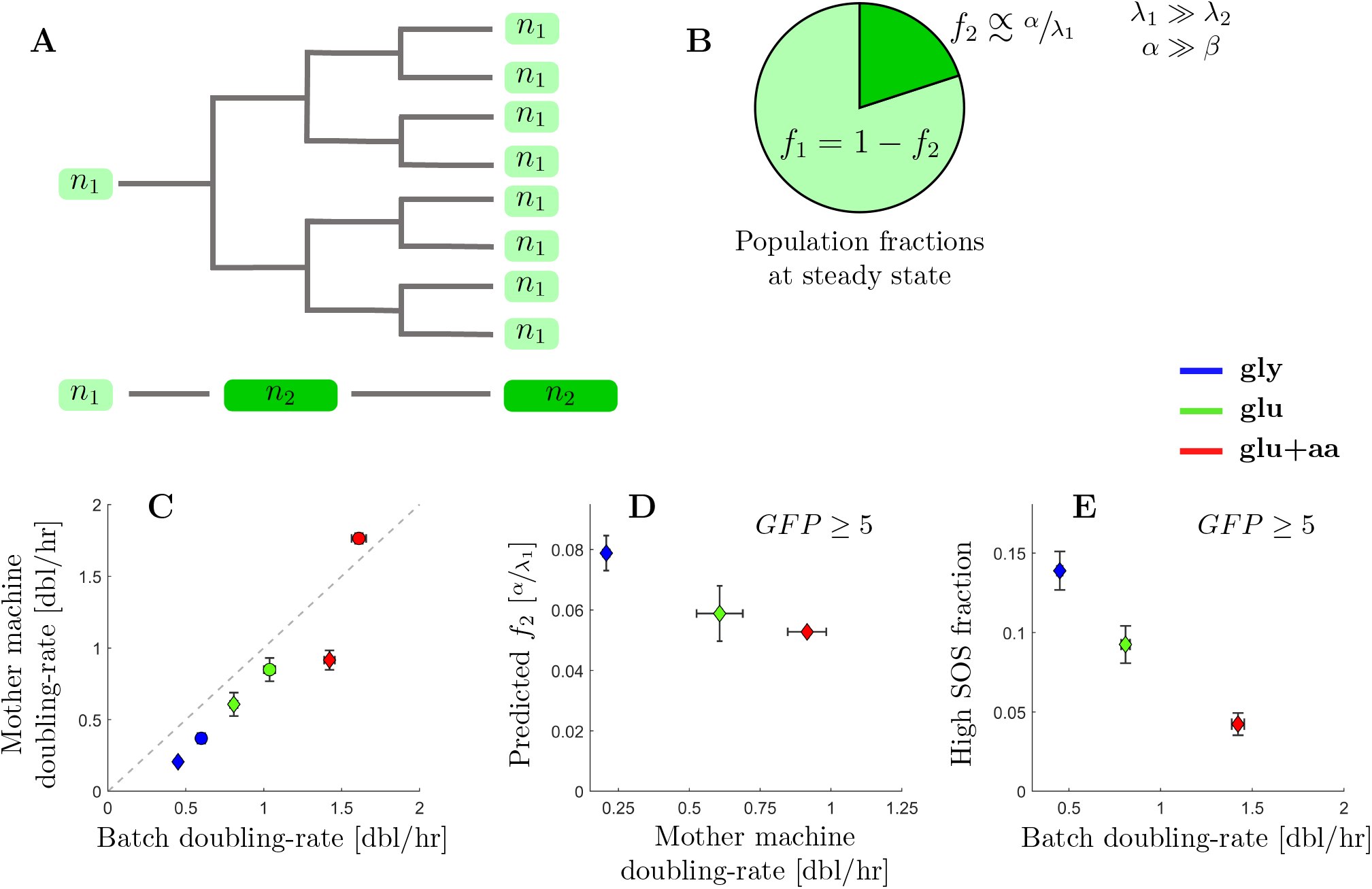
A two-populations model explains the large fraction of high SOS cells observed in slow growth conditions. A) Population model: Cells with high levels of SOS induction slow cell division and are outcompeted, in terms of cell numbers, by lineages experiencing low levels of SOS induction. B) In conditions when rate of switching to high SOS is much higher that the reverse and where the high SOS cells divide very slowly, the population fraction of cells with high SOS levels is expected to be inversely proportional to the growth-rate of the subpopulation with low SOS levels (at steady-state). C) Comparison between doubling rates for the WT (circles) and 2-pal mutant (diamonds) estimated from growth in the mother machine and in batch in different growth conditions. Mother machine doubling rates are the estimated population doubling rate derived from the underlying single-cell division-rate distribution. Points represent the average and bars the standard error from biological repeats. D) Predicted steady-state population fractions for high SOS cells (with a threshold of 5 arbitrary units) from the SOS reporter *PsulA-mGFP* under replication-dependent DSBs (2-pal) in different growth conditions. Points represent the average and bars the standard error from three biological repeats. The data are the prediction using rates estimated from mother machine experiments. E) Batch population fractions of high SOS cells (above 5 arbitrary units) from the SOS reporter *PsulA-mGFP* under replication-dependent DSBs (2-pal) in different growth conditions. Points represent the average and bars the standard error from three biological repeats.

When computing the expected fraction of high SOS cells (*f*_2_), we observed that our model indeed predicts a higher fraction of high SOS cells in poor nutrient conditions (despite lower rate of SOS induction) than in rich media (Figure 4D). For example, the expected fraction of cells reaching a GFP fluorescent intensity of at least 5 A.U. is 7.81±0.6% in M9-gly, 5.92±0.9% in M9-glu and 5.25±0.1% in M9-glu+aa. This is similar to the trend we measured in batch experiments (Figure 4E, respectively 14±1% in M9-gly, 9±1% in M9-glu and 4.3±0.7% in M9-glu+aa). However, our model tends to underestimate the faction of high SOS cells especially in low nutrient conditions. For example, we observe 14±1% cells above 5 A.U. versus a prediction of 7.81±0.6% in M9-gly. This might be explained by the generally lower division rate we observe in the mother machine set-up which may lead to lower rate of SOS induction in this set-up. Indeed, we expect DSBs arising from the presence of palindromes on the chromosome to correlate with replication rate and therefore division rates (Eykelenboom *et al*, 2008; Cockram *et al*, 2015; Amarh *et al*, 2018). Thus our model explains the counter-intuitive result that high SOS cells are more frequent in poor nutrient conditions despite the high rate of SOS induction in fast growth condition.

## DISCUSSION

In natural environments, bacteria are exposed to varying levels of nutrient availability and subject to sub-lethal stresses (Andersson & Hughes, 2014). The induction of stress responses, such the DNA damage response (SOS), consequently plays an important role in survival. In this paper, we show that heterogeneity in the levels of SOS expression induced by chronic sub-lethal DNA-damage exhibits strong growth-dependence. Surprisingly, we observe a larger fraction of highly induced cells in poor nutrient conditions despite a higher rate of SOS induction in rich nutrient conditions. This counter-intuitive result can be explained by the dynamic balance between the rate of SOS induction and the attenuated division rates for high-SOS induced cells.

At the single-cell level, SOS induction decouples growth from division. Our results indicate that this decoupling has a major impact on the dynamics of the population. Classically, during balanced growth the population doubling rate (µ) is directly related to the exponential rate of mass accumulation (λ=ln2*µ) (Monod, 1949). When cells induce the SOS response, they become filamentous because they continue adding mass whilst delaying division as a result of the induction SulA, YmfM and potentially other division inhibitors (Ansari *et al*, 2021). Heterogeneity in the level of SOS induction produces strong heterogeneity in the division rates of individual cells in the population. As we have shown, this leads to counter-intuitive effects at the population level, where the frequency of high SOS cells is not simply the result of the rate of SOS induction but also depends upon competition with the rapidly-proliferating non-induced cells.

Our work highlights the importance of a multi-scale approach to the analysis of bacterial stress responses. In the case of sub-lethal DNA damage, analysis of the population-averaged SOS induction level shows a negative growth-dependence, with higher induction in poor-nutrient conditions than in rich-nutrient ones (Supplementary Figure 5 A, B). A negative correlation between expression level and growth-rate is consistent with the growth-dependence of a fully-induced protein, and could suggest that rich nutrient conditions impose limits on SOS induction in order to express the requisite translation machinery (Hui *et al*, 2015; Scott *et al*, 2010; Weiße *et al*, 2015). Analysis at the single cell level, however, shows a somewhat different picture. For the majority of the population (bottom 85%, Figure 2C, D) we do not observe any growth dependence of the expression level of SOS genes. This could be explained by the negative auto-regulation of LexA at low level of SOS induction; negative feedback results in homeostatic expression levels, abrogating any intrinsic growth-rate dependence (Klumpp *et al*, 2009). For the high SOS cells (top 15%, Figure 2E, F), we do observe an average negative growth-dependence in the expression level of SOS genes, consistent with physiological constraints characterized in balanced growth. The dynamic equilibrium of these phenotypically-distinct subpopulations is maintained by a balance between the rate of SOS induction (α in our model) and the growth rates of the two subpopulations. Though the growth-dependence of the expression of SOS genes in each subpopulation conforms to what is known from physiological constraints on gene expression, our understanding of the population behavior critically depends upon quantification of the dynamics of molecular processes at the single-cell level via the SOS induction rate α.

The induction of the SOS response is known to have multiple consequences beyond facilitating the repair of DNA (Podlesek & Žgur Bertok, 2020), including increasing antibiotic tolerance (Dörr *et al*, 2009; Wu *et al*, 2015), modulating expression of mobile genetic elements (Beaber *et al*, 2004; Baharoglu *et al*, 2010; Fornelos *et al*, 2016), and increasing the rate of mutagenesis (Vaisman *et al*, 2012; Dapa *et al*, 2017; Pribis *et al*, 2019). Our observed growth-dependent heterogeneity in the fraction of SOS-induced cells suggests that care must be taken when making quantitative estimates of the mutation rate under conditions of sub-lethal DNA damage. For example, if the majority of mutants generated in a fluctuation assay arise from the high-SOS fraction, then the inferred mutation rate must be corrected (via multiplication by the reciprocal fraction of high-SOS cells, 1/f2) to account for the small subpopulation size. Quantitative predictions of the mutation rate in SOS-induced cells could therefore be underestimated by a factor of 20 or more. Furthermore, the correction to subpopulation size is growth-rate dependent (Figure 4E), introducing an inherent growth-rate dependence in the mutation rate. Given the growth-dependent heterogeneity in the population under SOS-induction, microfluidic single-cell mutation-accumulation assays (Uphoff, 2018; Robert *et al*, 2018) offer a useful tool to deconvolve heterogeneous stress response from downstream genetic change.

We have shown that single cell heterogeneity in division times can have important consequences in the abundance of cells with high SOS levels in growing populations. Similar heterogeneous population dynamics have been used to describe the maintenance of persister fractions (Balaban *et al*, 2004; Patra & Klumpp, 2013), and to describe phase-transitions in the stability of antibiotic-resistant strains (Roy & Klumpp, 2018; Deris *et al*, 2013). In addition to DNA damaging antibiotics, a growth-dependent heterogeneous response is likely to occur in treatments with cell-wall targeting antibiotics, as they can inhibit cell division, induce moderate levels of the SOS response, and induce the general stress response (Lambert & Kussell, 2015; Laureti *et al*, 2013; Miller *et al*, 2003). More generally, our results argue that whenever a stress leads to a transition towards a non-dividing or slow-dividing state (in our case high SOS expression), the fraction of these cells will be enriched in slow-growth conditions. This is likely to affect the fraction of subpopulations in natural environments with varying levels of nutrient availability, and introduces an intrinsic growth-rate dependence in bet-hedging strategies (Veening *et al*, 2008). Growth-rate dependent heterogeneity under DNA-damaging conditions introduces an additional degree of freedom in the complex coupling between the growth environment and evolutionary change.

## MATERIALS AND METHODS

### Culture conditions

For all microscopy and batch experiments cell cultures were grown in M9 based media. The composition of the M9 salts was: 49 mM Na2HPO4, 22 mM KH2PO4, 8.6 mM NaCl, 19 mM NH4Cl, 2 mM MgSO4, and 0.1 mM CaCl2. This was supplemented with either 0.5% w/v glycerol or 0.5% w/w glucose, and a mix of amino-acids (1X MEM Non-Essential Amino Acids and 1X MEM Amino Acids, both manufactured by Gibco R). For strains and plasmid construction cells were grown in LB, or LB agar supplemented with the corresponding selection markers. Concentrations employed for antibiotics were: ampicillin 100 µg/ml, kanamycin 50 µg/ml, and gentamycin 10 µg/ml. All cultures were grown in 50 ml falcon tubes agitated at 37°C (300 rpm) with no more than 5 ml of liquid volume, unless otherwise stated.

Cell cultures were grown for at least 12 division times in each media to reach steady-state exponential growth before taking measurements. This was carried out as follows. Cells were taken from frozen stocks at −80°C, and grown for 10-16 hours in LB media. They were then grown overnight (after a 1:1000 dilution) in their respective M9-based media, plus appropriate antibiotics in case of a selection marker. These overnight cultures were diluted 1:200 in fresh media (without antibiotic), and grown until OD600 0.1 (approximately three division times). From there, they were diluted again in fresh media (with dilution in the order of 10^−5^ to 10^−6^) so that experiments could be performed the next day. These last dilution factors were calculated to allow for at least 12 division times (without reaching an OD600 higher than 0.15) before any measurement. During the day, samples were diluted when necessary in order to prevent reaching OD600 higher than 0.15. Population (batch) growth rates were estimated by OD600 measurements over time in three technical repeats and three biological repeats per condition.

### Strain and plasmid construction

*E. coli* MG1655 was used as WT strain in this study. The strains and plasmids are listed in Table 1 and 2 of the supplementary material. Gene-expression reporters (GFP for SOS expression and mKate for constitutive gene expression) were cloned into pOSIP plasmids inserted into the genome by clone-integration (St-Pierre *et al*, 2013). Plasmid construction was performed by Gibson assembly after PCR amplification of the fragments (see Table 3 in the supplementary material for the detailed description and Table 4 for the list of primers used). All strains were checked with PCR amplification followed by Sanger sequencing. Insertion of interrupted palindromes performed via P1 transduction using strains kindly given by D. L. Leach (see Table 1 in supplementary material).

### Fluorescence microscopy

All images were captured using a Nikon Ti-E inverted microscope equipped with EMCCD Camera (iXion Ultra 897, Andor), a SpectraX Line engine (Lumencor) and a 100X Nikon TIRF objective (NA 1.49, oil immersion). Nikon Perfect-Focus system was used for continuous maintenance of focus. The filter set for imaging *mGFP* consisted of ET480/40x (excitation), T510LPXR (dichroic), and ET535/50m (emission); whereas for *mKate2* the set ET572/35x (excitation), T590LPXR (dichroic), and ET632/60m (emission) was used. Filters used were purchased from Chroma. GFP fluorescence was measured using 80 milliseconds exposure, whereas mKate2 fluorescence was imaged for 100 milliseconds, both at minimal gain and maximum lamp intensity. Microscope was controlled from MATLAB via MicroManager (Edelstein *et al*, 2014).

### Agar-pad microscopy and image analysis

For agar-pad microscopy, steady-state exponential cell cultures were prepared as described previously. For imaging samples were mounted on agar-pads: 5-10 µl from cultures around OD600 0.05 were placed in 1% agarose pads (Gene-frame 65 µl) made with the corresponding growth media. For each repeat, about 250 stage positions were imaged, comprising a total of 4000-30000 cells after image analysis. All conditions were performed in at least three biological repeats. For time-lapse microscopy experiments, 2 µl from liquid cultures at balanced growth were placed in agar-pads as described above. About 15 different stages positions were imaged at intervals lasting one tenth of the population doubling time, for one hundred time intervals.

In order to automate the detection of cells from fluorescent images, we developed an algorithm for edge-detection using custom low-pass filters (the algorithm is detailed in the supplementary material). Results from the automated cell segmentation were manually curated to remove any misidentified cell and false positives. Fluorescence signal from the constitutive reporter *PtetO-mKate2* was used in all cases for cell segmentation. All *mGFP* and *mKate* fluorescence values were re-scaled by the average fluorescence value of the WT strain data sets in each growth medium. After cell segmentation, fluorescence signal concentration was quantified by summing the total intensity for each cell divided by the number of pixels. For the population frequency plots, data were binned and plotted in log-scale intervals (the mean ± standard error across biological repeats is reported).

### Mother machine experiments

#### Microfluidics design and fabrication

We used a mother machine design similar to (Wang *et al*, 2010), consisting of an array of closed-end microchannels connected to a large flow channel. The device was designed using OpenSCAD and the photomask was manufactured by Compugraphics International Ltd. The master moulds were produced at the Scottish Microelectronics Centre, Edinburgh, using standard soft lithography techniques and SU-8 photoresists on a 4” silicon wafer. This was done in two steps: the first layer for the microchannels and the second layer for the flow channel. The length of the microchannels were ∼25 µm and the height of the flow channel was ∼22 µm. The size of *E. coli* cells change depending on the medium used -this scales with growth rate. To accommodate for this, our design consisted of microchannels with a range of widths (from 0.9-1.9 µm) and several master moulds were fabricated corresponding to different heights (0.9-1.36 µm). Appropriate dimensions were tested and selected for each growth condition. See Table 5 in the supplementary material for specific dimensions used. The poly-dimethylsiloxane (PDMS) chips were made using the Silicone Elastomer Kit 184 (Sylgard, Dow Corning) with a 1:10 ratio of curing agent to base. The protocol used to fabricate the microfluidics chips is summarized in the supplementary material.

#### Culture preparation

10 ml of cultures were grown into steady-state exponential phase using similar pre-culture conditions described earlier. Cells were harvested at OD600≈0.2 and concentrated 100-fold by centrifugation (4000 rpm for 5 min). Tween-20 (Thermo Scientific Pierce) was added to the culture (0.01% final concentration) before centrifugation to prevent clumping. Before sample loading, the chip was passivated with Tween-20 (0.01%) for at least 1 h. The concentrated cell culture was then injected into the feeding channels using a 1 ml syringe with a 21-guage blunt needle (OctoInkjet). Cells were allowed to diffuse into the microchannels for approximately 30 min at 37°C. To further assist loading, the cells were then spun into the microchannels by centrifugation at 3220 ×g for 5 min using a custom built mount. The microfluidics device was then mounted on the microscope and connected to a peristaltic pump (Ismatec IPC ISM932D) on one end, and to fresh media + 0.01% Tween-20 on the other end, in order to flow fresh media through the device. Cells were flushed from the main trench at 1.5-2 ml/h and then the flow rate lowered to 1 ml/h for the duration of the experiment. Experiments were run for 10-42 hours depending on the nutrient conditions. Cells in the channels were allowed to recover for at least 2 h at 37°C before imaging.

#### Microscopy and image analysis

For mother machine experiments, images were acquired at 5 min intervals for M9-glu+aa, 10 min for M9-glu, and 12 min for M9-gly. Images were saved in .mat format as one file per fluorescence channel per frame. The images were then converted to TIFF format in MATLAB. Segmentation and tracking was performed using BACMMAN run in Fiji (Ollion *et al*, 2019). The BACMMAN configuration was adapted to segment and track cells based on fluorescence images. Images were imported in BACMMAN and pre-processed prior to segmentation. This included image rotation to ensure channels were vertically oriented and the channel opening was at the bottom, and cropping of images to include only the area consisting of microchannels. Joint segmentation and tracking of cells were performed on the *mKate2* fluorescence channel. Segmentation parameters were optimised for each data set. Curation of segmentation and tracking was carried out using BACMMAN’s interactive graphical interface. Although automated segmentation and tracking was mostly accurate, occasionally errors were produced. Thus, every lineage was manually checked, with 2-pal data sets requiring the most curation mainly due to excessive elongation of cells expressing the SOS response. Mother cells that did not grow for the duration of the experiment and those that were already excessively elongated at the beginning of the experiment were removed. Positions with channel deformities and where loss of focus occurred were also discarded. Cell fluorescence and morphology variables were then exported from BACMMAN into excel files for further processing.

Further analysis was performed using custom MATLAB scripts. First, data corresponding to the ‘mother cell’ lineage was isolated. Divisions for the mother cell lineage were recorded with tracking continuing for the cell at the closed end of the channel and discarding the sister cell. Division rates were calculated as the inverse of the interdivision time. The population growth rate was then estimated from the division rates as explained in the supplementary material. Cell elongation rates were calculated using a linear fit to the logarithm of cell length as a function of time per cell cycle. Cell length was determined as the maximal distance between two points of the cell contour as reported by BACMMAN. A minimum of 3 data points per generation was imposed as a fitting constraint and negative growth rates were removed. The rate of reaching a high SOS state (α) was estimated based on GFP fluorescent value as described in the supplementary material.

### Data availability

The models and algorithms developed in this study are described in the supplementary methods.

The microscopy images will be deposited on the open access Datashare server of hte University of Edinburgh and will be accessible through a doi.

The parameters estimated from image analysis and used as parameters for the models are summarizes in supplementary Table 1 (snapshot data) and Table (Mother Machine data).

## Supporting information

Supplementary data: Agar-pad snapshot values

Supplementary data: Mother machine values

## Acknowledgements

We wish to thank Karolina Gaebe and Aleksandra Jarseva for performing preliminary characterization, and Nate Lord for supplying the strain used to amplify the *mKate2* gene. We are grateful to David Leach and Benura Azeroglu for helpful discussions. We wish to thank Juan Carlos Arias Castro, Pascal Hersen, Dario Miroli, Eric Thorand and Filippo Menolascina for useful advice and discussions on the usage and design of the mother machine and microfluidics. The master production was performed in the Scottish Microelectronics Centre (SMC), the School of Engineering’s small research facility, with the help of SMC’s staff. This work has been supported by a Wellcome Trust Investigator Award 205008/Z/16/Z (to M.E.K.), a Natural Sciences and Engineering Research Council of Canada (NSERC) Discovery grant (to MS), the UKRI Synthetic Biology for Growth programme and BBSRC/EPSRC/MRC funded Synthetic Biology Research Centre BB/M018040/1 (to T.P.), and two Darwin Trust of Edinburgh postgraduate studentships (to S.J.R and J.B.).

## Author Contributions

S.J.R., M.S, and M.E.K. conceived the experiments. S.J.R. and J.B. performed the experiments. A.MV, S.J.R, T.P. and M.E.K. developed and A.MV and S.J.R designed microfluidic technology. S.J.R, J.B. and M.E.K. analyzed the data. S.J.R., J.B., M.S, and M.E.K. interpreted the data and wrote the manuscript. All authors reviewed the manuscript.

## Supplementary Figures

**Supplementary figure 1:**
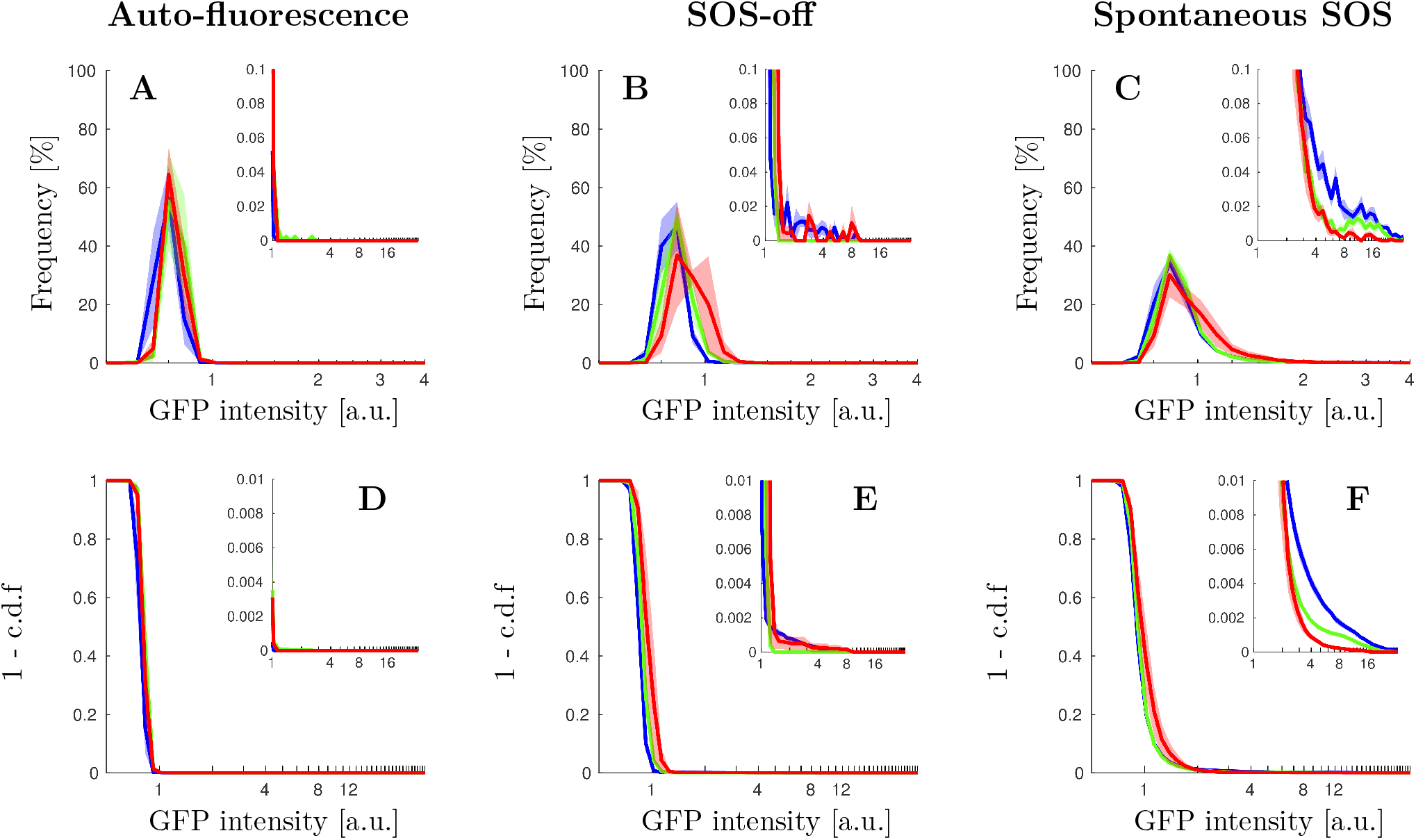
Single cell distributions of GFP intensity without inducing DNA damage. For all plots, growth conditions are: M9-glycerol (blue), M9-glucose (green), and M9-glucose+amino-acids (red). Solid lines represent the average and shaded area the standard error from biological repeats. A) Steady state distribution of GFP intensity from cell auto-fluorescence in different growth conditions. B) Steady state distribution of GFP intensity from SOS reporter *PsulA-mGFP* for cells unable to induce SOS (SOS-off, *lexA3* background) in different growth conditions. C) Steady state distribution of GFP intensity from SOS reporter *PsulA-mGFP* for wild type cells in different growth conditions. D) Steady state cumulative distribution of GFP intensity from cell auto-fluorescence in different growth conditions. E) Steady state cumulative distribution of GFP intensity from SOS reporter *PsulA-mGFP* for cells unable to induce SOS (SOS-off, *lexA3* background) in different growth conditions. F) Steady state distribution of GFP intensity from SOS reporter *PsulA-mGFP* for wild type cells in different growth conditions.

**Supplementary figure 2:**
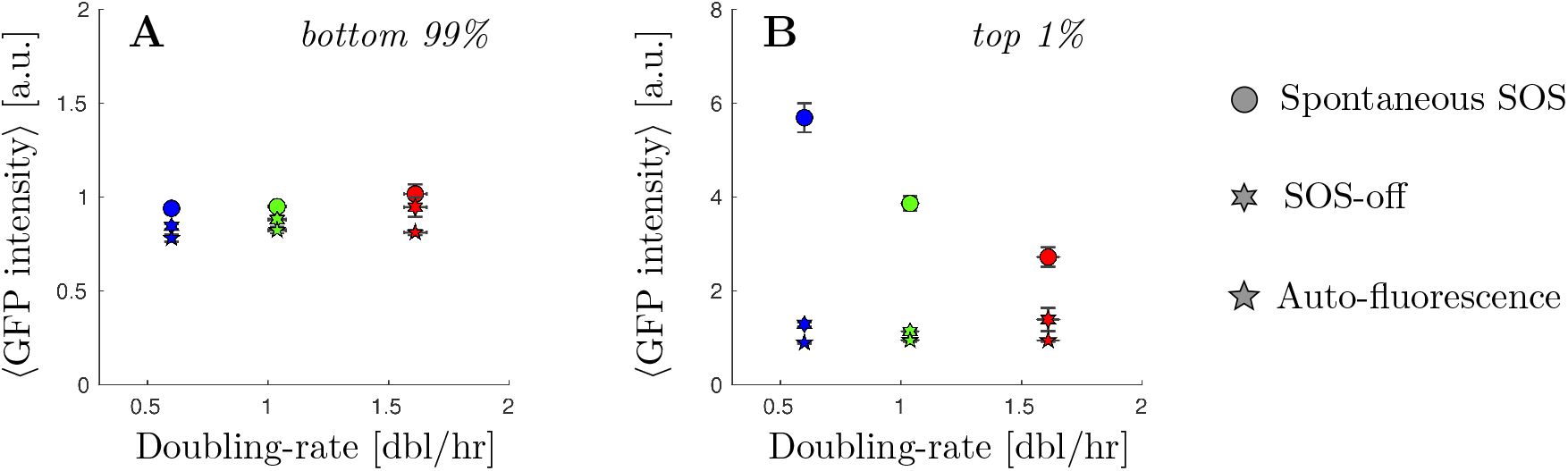
Average GFP intensity without inducing DNA damage. For all plots, growth conditions are: M9-glycerol (blue), M9-glucose (green), and M9-glucose+amino-acids (red). Points (dots WT strain, 6-points-stars SOS-off, *lexA3* background, 4-point-stars, autofluorescence) represent the average and bars the standard error from biological repeats. A) Average GFP intensity for the bottom 99% of the population in different growth conditions. B) Average GFP intensity for the top 1% of the population in different growth conditions.

**Supplementary figure 3:**
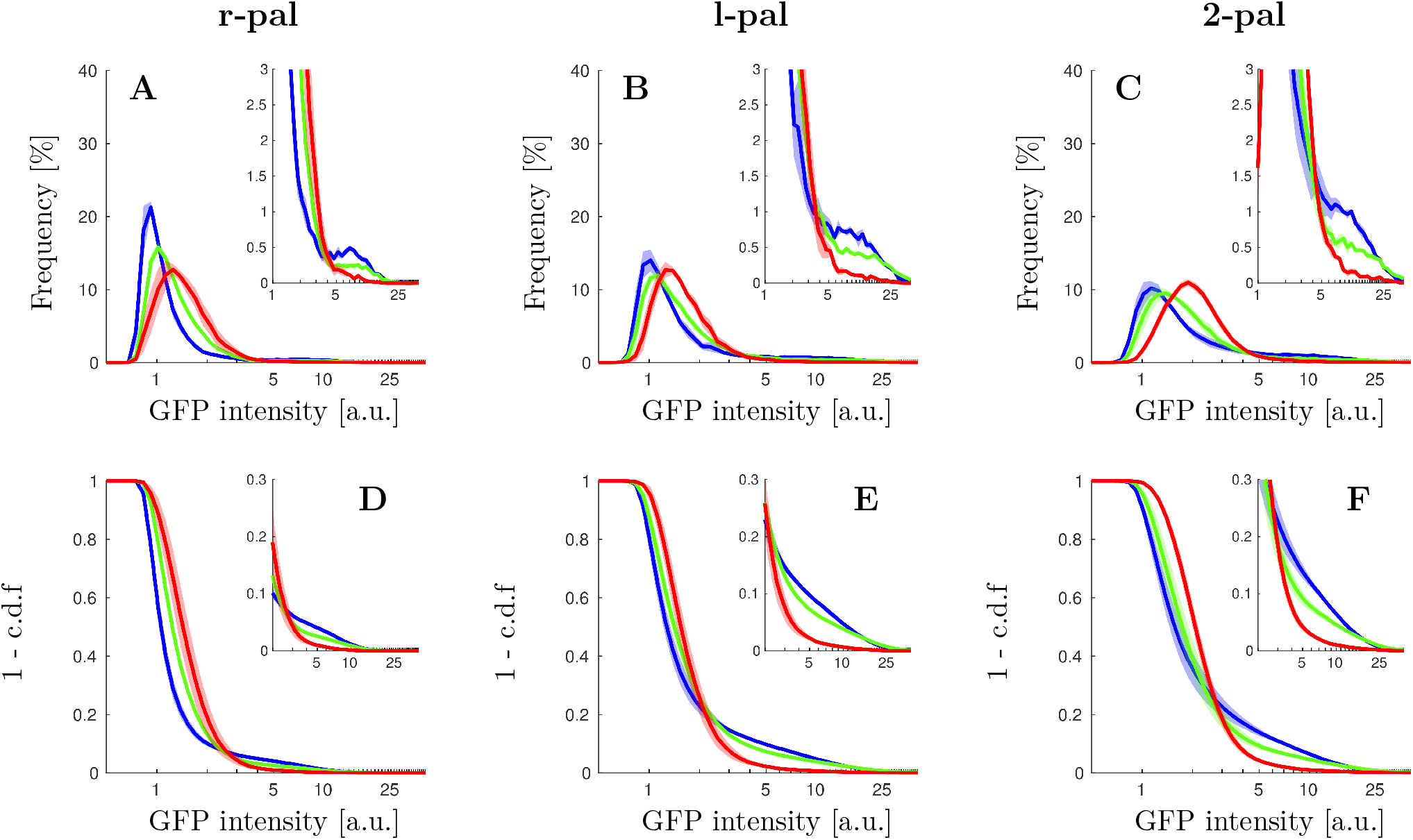
Single cell distributions of GFP intensity under replication-dependent DSBs. For all plots, growth conditions are: M9-glycerol (blue), M9-glucose (green), and M9-glucose+amino-acids (red). Solid lines represent the average and shaded area the standard error from at least 3 replicates done in different days. A) Steady state distribution of GFP intensity from SOS reporter *PsulA-mGFP* for cells under replication-dependent DSBs from the r-pal palindrome, in different growth conditions. B) Steady state distribution of GFP intensity from SOS reporter *PsulA-mGFP* for cells under replication-dependent DSBs from the l-pal palindrome, in different growth conditions. C) Steady state distribution of GFP intensity from SOS reporter *PsulA-mGFP* for cells under replication-dependent DSBs from both l-pal and r-pal palindromes (2-pal), in different growth conditions. D) Steady state cumulative distribution of GFP intensity from SOS reporter *PsulA-mGFP* for cells under replication-dependent DSBs from the r-pal palindrome, in different growth conditions. E) Steady state cumulative distribution of GFP intensity from SOS reporter *PsulA-mGFP* for cells under replication-dependent DSBs from the l-pal palindrome, in different growth conditions. F) Steady state cumulative distribution of GFP intensity from SOS reporter *PsulA-mGFP* for cells under replication-dependent DSBs from both l-pal and r-pal palindromes (2-pal), in different growth conditions.

**Supplementary figure 4:**
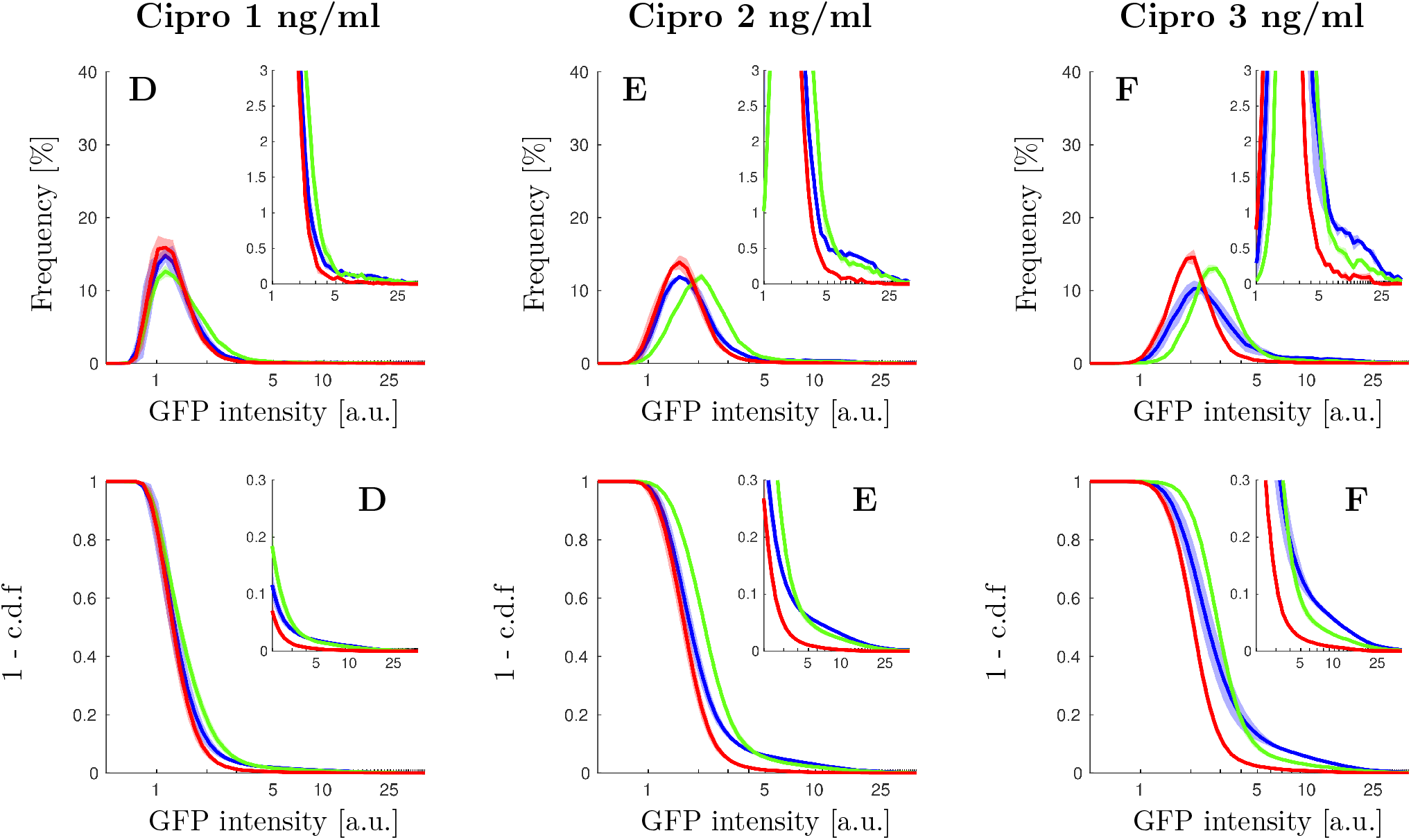
Single cell distributions of GFP intensity under ciprofloxacin. For all plots, growth conditions are: M9-glycerol (blue), M9-glucose (green), and M9-glucose+amino-acids (red). Solid lines represent the average and shaded area the standard error from at least 3 replicates done in different days. A,B,C) Steady state distribution of GFP intensity from SOS reporter *PsulA-mGFP* for cells exposed to 1 ng/ml (respectively 2 ng/ml, 3 ng/ml) of ciprofloxacin in different growth conditions. D,E,F) Steady state cumulative distribution of GFP intensity from SOS reporter *PsulA-mGFP* for cells exposed to 1 ng/ml (respectively 2 ng/ml, 3 ng/ml) of ciprofloxacin in different growth conditions.

**Supplementary figure 5:**
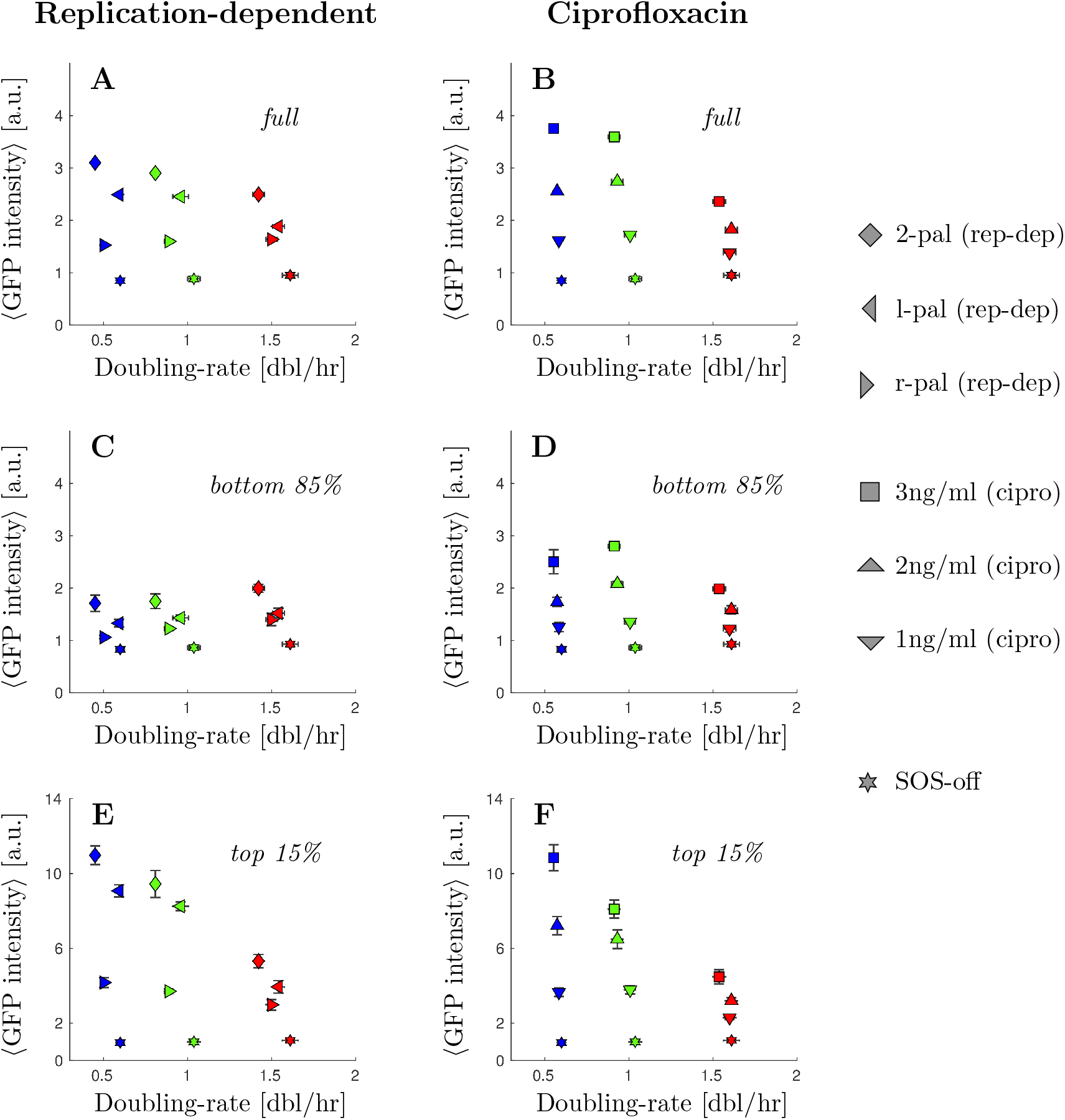
Average GFP intensity from inducing double-strand breaks. For all plots, growth conditions are: M9-glycerol (blue), M9-glucose (green), and M9-glucose+amino-acids (red). Points represent the average and bars the standard error from at least 3 replicates done in different days. A) Average GFP intensity for the whole population under replication-dependent DSBs as a function of growth rate. B) Average GFP intensity for the whole population under ciprofloxacin as a function of growth rate. C) Average GFP intensity for the bottom 85% of the population under replication-dependent DSBs as a function of growth rate. D) Average GFP intensity for the bottom 85% of the population under ciprofloxacin as a function of growth rate. E) Average GFP intensity for the top 1% of the population under replication-dependent DSBs as a function of growth rate. F) Average GFP intensity for the top 1% of the population under ciprofloxacin as a function of growth rate.

**Supplementary figure 6:**
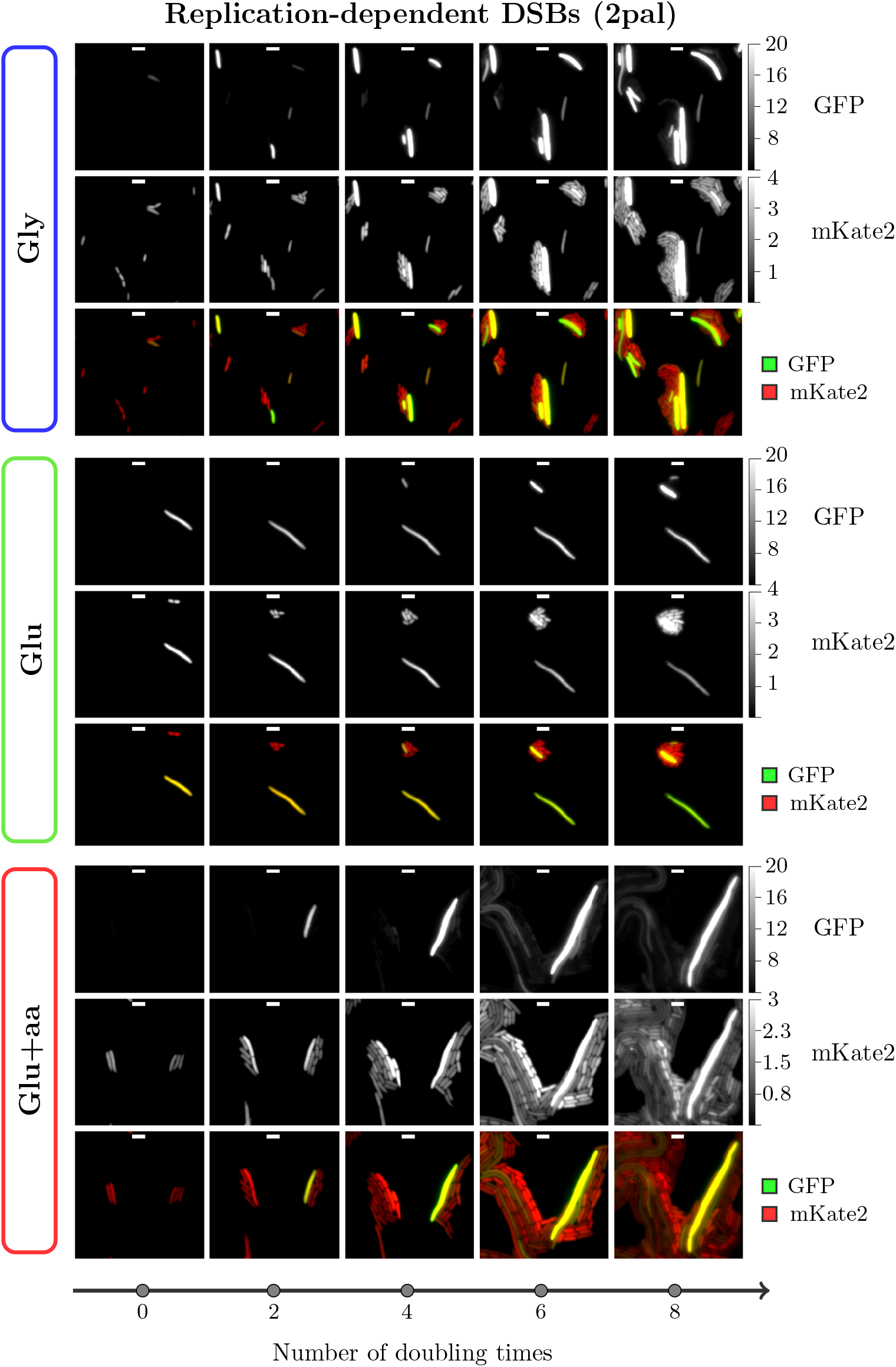
Time-lapse microscopy of cells under replication-dependent DSBs. Cells carrying two palindromes, the SOS reporter *PsulA-mGFP*, and a constitutively expressed reporter *PtetO-mKate2*, were imaged using agar-pads made with different growth media. Cells were grown to steady-state exponential growth before mounting into the agar-pads.

**Supplementary figure 7:**
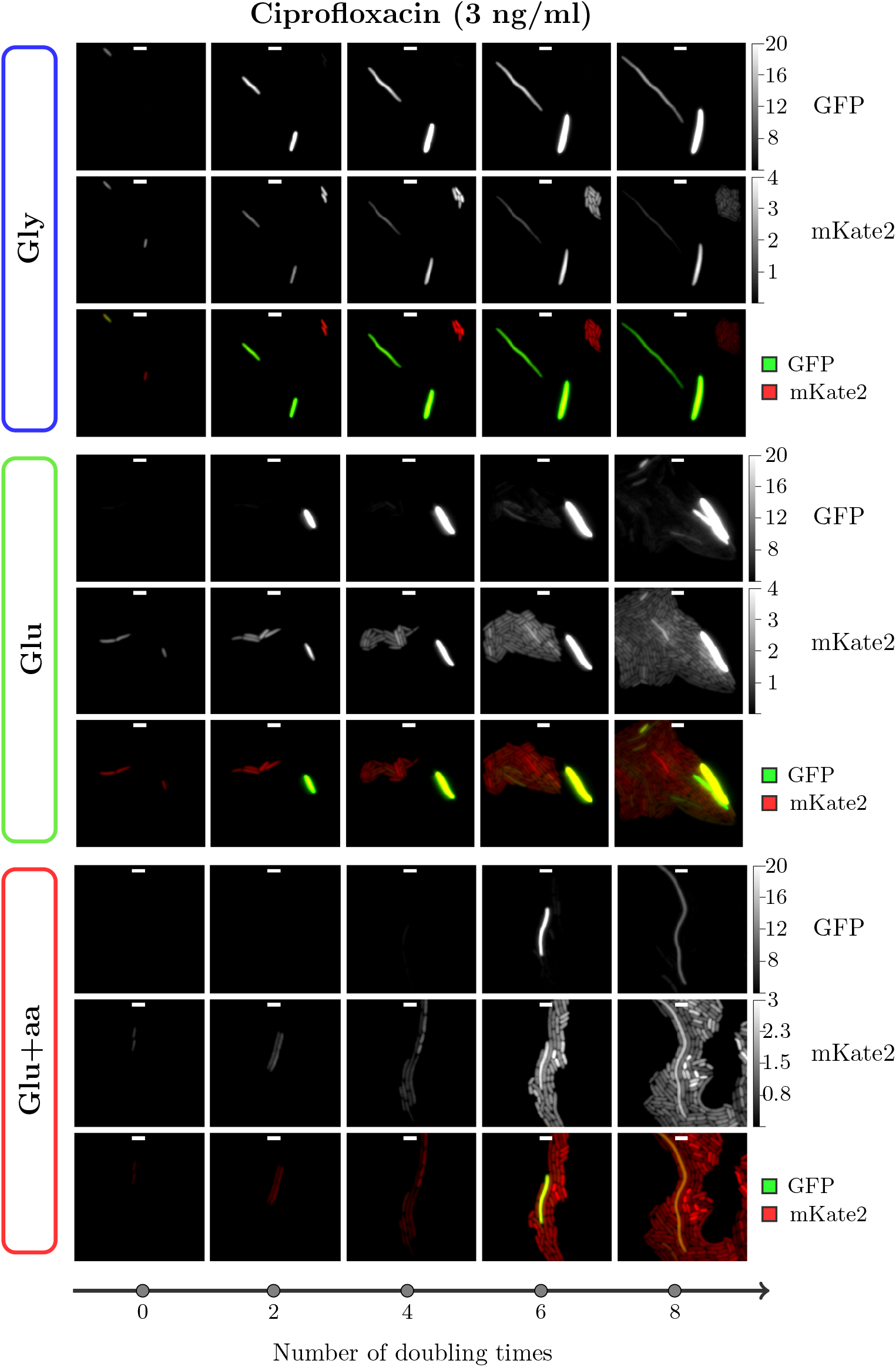
Time-lapse microscopy of cells exposed to ciprofloxacin. Cells carrying the SOS reporter *PsulA-mGFP*, and a constitutively expressed reporter *PtetO-mKate2*, were imaged using agar-pads made with different growth media supplemented with 3 ng/ml of ciprofloxacin. Cells were grown to steady-state exponential growth in exposure to the antibiotic before mounting into the agar-pads.

**Supplementary figure 8:**
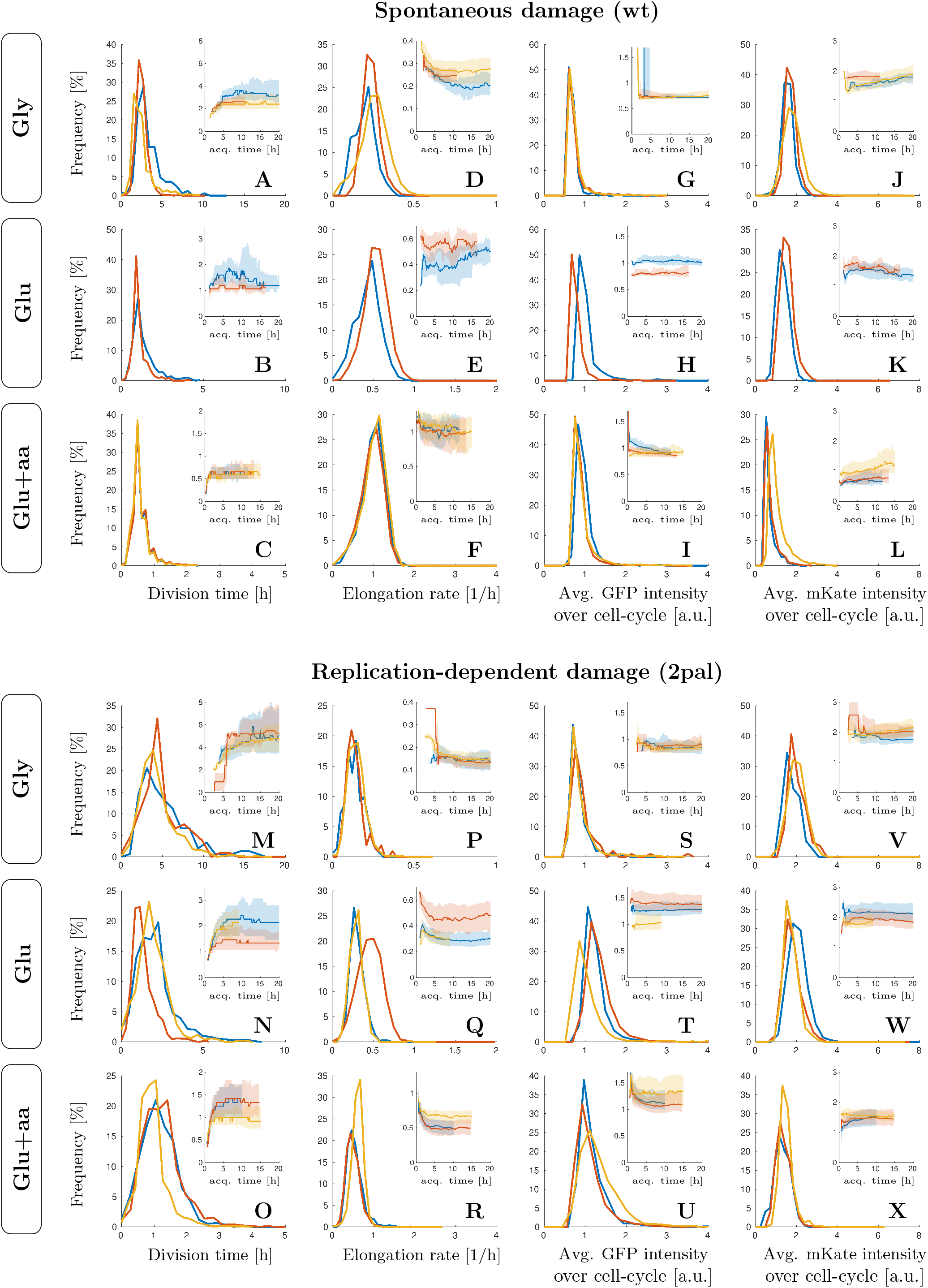
Distribution of main single-cell parameters from mother-machine experiments. The lineages of wild-type cells and cells undergoing replication dependent DNA-damage were tracked for each cell cycle, and their division time, elongation rate, average fluorescence intensity in the GFP and mKate2 channels recorded. The distribution of these 4 values are presented. Each color (Red, yellow and blue) represents an independent biological repeat. On each panel inset, we show the median values over time, and with shaded areas representing the first and third quartile of each distribution. For each timepoint in the inset curves, only cell-cycles spanning that particular time-point were included for computations of the median, first and second quartiles.

**Supplementary figure 9:**
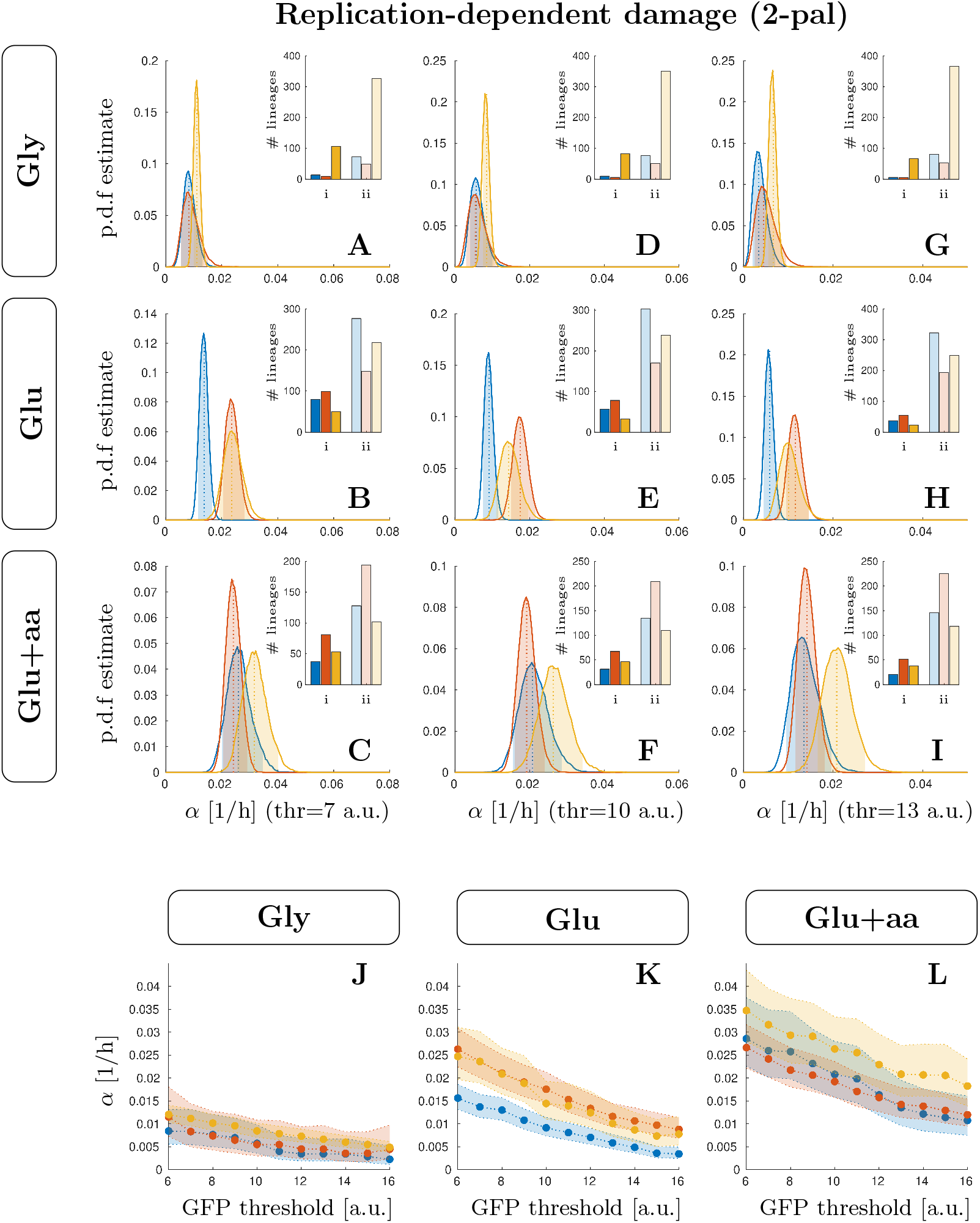
Inference of the transition rate to high SOS from individual cell traces at different GFP intensity thresholds. The single cell GFP intensity trajectories were used to estimate the transition rate constant (*α*) based on different GFP intensity threshold. The posterior probability of based on the observations was computed from dividing lineages into those that cross and do no cross the threshold, those are case “i” and “ii” respectively. Examples of these posterior probabilities are presented in panels A-I, where the shared areas represent the range between the 5th and 95th percentiles. The inlets in panels A-I are used to show the number of lineages on each category. In panels J-L we show the most likely value of *α* for the different growth-conditions, and shaded areas represent the 5th and 95th percentiles from the posterior distributions. In all panels, the color red, yellow, and blue are used to represent three independent biological repeats.

