## Supplementary methods for "Growth-dependent heterogeneity in the DNA damage response in *Escherichia coli*"

May 5, 2021

#### Contents

**1 List of plasmids, strains, and primers** **2**

**2 Image analysis of snapshot data** **4**

**3 Mother machine** **6**
3.1 Microfluidics device fabrication . . . . . . . . . . . . . . . . . . . . . . . . . . . . . 6
3.2 Mother machine data analysis . . . . . . . . . . . . . . . . . . . . . . . . . . . . . 6

**4 Estimating the transition rate to the high SOS state from mother machine experiments** **7**

**5 Estimating the steady-state population growth-rate from single-cell division time distributions** **8**

**6 Estimating the fraction of high SOS cell in a growing population** **11**

### 1 List of plasmids, strains, and primers

Here are the list of strains, plasmids and primers used in this study. Bacterial strains were constructed either by P1 transduction or by cloneintegration (St-Pierre et al., 2013) as mentioned in Table 1. Plasmids pSJR036 and pSJR046 were constructed by digestion of the backbone by enzymatic restriction, amplification of the insert by PCR using a high fidelity polymerase and ligation using Gibson assembly (details in Table 3). All plasmids were checked by PCR amplification of the insert and Sanger sequencing. After construction, all strains were checked by PCR amplification and Sanger sequencing of the modified chromosomal region.

Table 1: **List of strains**. CLI stands for clone-integration, and P1 for phage transduction.

| **Strain** | **Background** | **Genotype** | **Source/Construction** |
|---|---|---|---|
| eSJR017 | MG1655 | *seqA::mGFP* $P_{rna1}$-*mKate2* | Gift from Raul Fernandez Lopez (RFL84). mKate2 construct built by Nathan Lord. |
| eSJR048 | MG1655 | *rph-1* $\lambda^-$ | Genomic Stock Center (CGSC7740) |
| eSJR059 | MG1655 | *lacI*$^q$ *lacZ::pal246 cynX::Gm*$^R$ | Gift from David Leach (DL2859) |
| eSJR130 | BW27784 | *asbB::pal246 ascF::Kn*$^R$ | Gift from David Leach (DL4212) |
| eSJR145 | MG1655 | *HK022:*$P_{sulA}$-*mGFP* | eSJR048 CLI using pSJR036 |
| eSJR206 | MG1655 | *HK022:*$P_{sulA}$-*mGFP* *P21:*$P_{tet01}$-*mKate2* | eSJR145 CLI using pSJR046 |
| eSJR214 | MG1655 | eSJR206 *asbB::pal246 ascF::Kn*$^R$ | eSJR206 P1 using eSJR130 |
| eSJR301 | MG1655 | eSJR206 *lacI*$^q$ *lacZ::pal246 cynX::Gm*$^R$ | eSJR206 P1 using eSJR059 |
| eSJR302 | MG1655 | eSJR206 *lacI*$^q$ *lacZ::pal246 cynX::Gm*$^R$ *asbB::pal246 ascF::Kn*$^R$ | eSJR301 P1 using eSJR130 |

Table 2: **List of plasmids**.

| **Plasmid** | **Purpose** | **Source** |
|---|---|---|
| pSJR017 | Clone-integration marker excision | pE-FLP (St-Pierre et al., 2013) |
| pSJR021 | Clone-integration at HK022 site | pOSIP-KH (St-Pierre et al., 2013) |
| pSJR025 | Clone-integration at P21 site | pOSIP-KT (St-Pierre et al., 2013) |
| pSJR035 | Source sequence for P*sulA-mGFP* | DL4847 |
| pSJR036 | $P_{sulA}$-*mGFP* insertion by CLI | This study |
| pSJR046 | $P_{tetO1}$-*mKate2* insertion by CLI | This study |

Table 3: **Plasmid construction**. List of plasmids constructed in this study.

| **Plasmid** | | **Backbone** | **Digestion** | **PCR template(s)** | **primer pair(s)** |
|---|---|---|---|---|---|
| pSJR036 | *HK022::*$\mathrm{P}_{sulA}$*-mGFP* | pSJR021 | EcoRI, PstI | pSJR035 | oSJR084, oSJR085 |
| pSJR046 | *P21::*$\mathrm{P}_{tetO1}$*-mKate2* | pSJR025 | EcoRI, PstI | eSJR017 | oSJR066, oSJR098 |

Table 4: **List of primers**.

| **Primer** | **5′-3′ Sequence** | **Purpose** |
|---|---|---|
| oSJR084 | GGACGCCCGCCATAAACTGCCAGGAATTGG GGATCGGAATTCAGGGTTGATCTTTGTTGT | pSJR036 construction |
| oSJR085 | TTAGGTTAGGCGCCATGCATCTCGAGGCAT GCCTGCAGTTATTTGTATAGTTCATCCATG | pSJR036 construction |
| oSJR066 | ACGCCCGCCATAAACTGCCAGGAATTGGGG ATCGGAATTCTTATCTGTGCCCCAGTTTGC | pSJR046 construction |
| oSJR098 | ATGAATTCAAATACTGTCCTTCCGGTCAGT GCGTCCTGCTGATGTGCTCAGTATCTCTAT CACTGATAGGGATGTCAATCTCTATCACTG ATAGGGACTCGACTGCAGGCATGCCTCGAG ATGCATGGCGCCTAACCTAAACTGACA | pSJR046 construction |
| oSJR058 | GGAATCAATGCCTGAGTG | HK022 insertion verification |
| oSJR059 | ACTTAACGGCTGACATGG | HK022 insertion verification |
| oSJR060 | ACGAGTATCGAGATGGCA | HK022 insertion verification |
| oSJR061 | GGCATCAACAGCACATTC | HK022 insertion verification |
| oSJR092 | ATCGCCTGTATGAACCTG | P21 insertion verification |
| oSJR093 | ACTTAACGGCTGACATGG | P21 insertion verification |
| oSJR094 | GGGAATTAATTCTTGAAGACG | P21 insertion verification |
| oSJR095 | TAGAACTACCACCTGACC | P21 insertion verification |
| oSJR072 | TTATGCTTCCGGCTCGTATG | *lacZ::pal246* verification FW |
| oSJR073 | GGCGATTAAGTTGGGTAACG | *lacZ::pal246* verification RV |
| oSJR080 | CCAACCAGTCTGAAGGTGCG | *ascB::pal246* verification FW |
| oSJR081 | CCAGCGGTTCGATACCGTAC | *ascB::pal246* verification RV |

#### 2 Image analysis of snapshot data

In order to automate the detection of cells from fluorescent images (cell segmentation), we developed a custom algorithm based on edge-detection using low-pass filters (detailed in Algorithm 1) and a graphical user interface to facilitate manual correction of the segmentation. The algorithm was designed to detect cell edges using a custom convolution filter, that compares each pixel value relative to its neighbours. The filters are constructed by summing many 2D Gaussian distributions, where each Gaussian has a mean position moving away from the center of the filter with a given orientation (detailed in Algorithm 2). We apply several filters with different orientations to the fluorescence image, and compute a score by combining the results from all filters (Algorithm 1). Then a threshold is applied to the score to remove cell edges, and generate a mask from which cells are identified as individual connected components (see example in Figure 1). Finally, the resulting segmentation is manually curated to remove any potential misidentified cell.

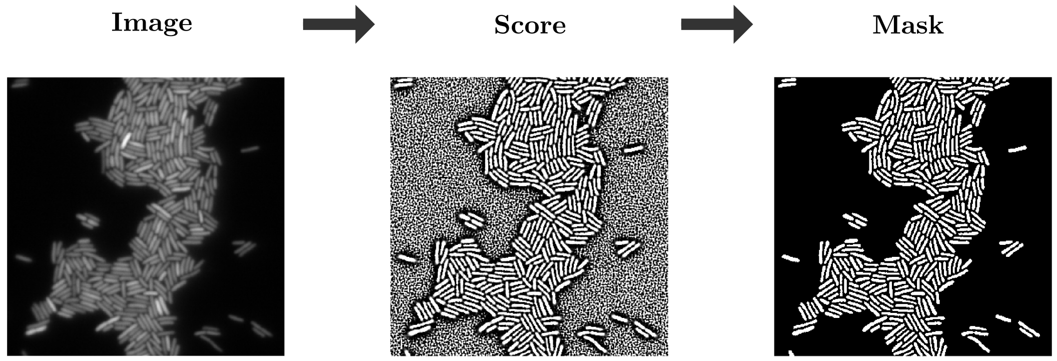


Figure 1: **Semi-automated cell detection**. Example of computed score and mask for a given image.

**Algorithm 1:** Cell segmentation from fluorescence image.
Algorithm for segmenting a fluorescent image using an array of spacial low-pass filters. It takes any image as input, plus seven parameters, and returns a mask containing where regions that appears as "valleys" in the intensity landscape have been removed

**Require:** Input image: $img$. Parameters: minimum intensity value $i_0$; $\mu$ and $\sigma$ (gaussian filter); pixel length $d$, width $w$, and set of angles $A = \{A_1, A_1 + \pi\}$ (for low pass filter); and a score threshold $s_0$. Some predefined functions: IMFILTER that applies a convolution filter to an image; GAUSSFILTER that return a gaussian filter; THRESHOLD that thresholds an image returning a boolean matrix; LOWPASSFILTERS that computes custom low-pass filters (see algorithm 2); IMCOMPLEMENT that computes the complement of an image; POSBOOL that returns one if the value is positive; PAIRWMULT that computes the pairwise multiplication of matrices; and PAIRWDIV that computes the pairwise division of matrices.

---

**function** SEGMENTATION_MASK($img$,$i_0$,$\mu$,$\sigma$,$d$,$w$,$A$,$s_0$) ▷ Returns $mask$
  $img \leftarrow$ IMFILTER(GAUSSFILTER($\mu, \sigma$), $img$) ▷ Filter image noise
  $mask0 \leftarrow$ THRESHOLD($img, i_0$) ▷ Threshold image
  $Filts \leftarrow$ LOWPASSFILTERS($d, w, A$) ▷ Set low pass filters
  $na \leftarrow$ LENGTH($A$) ▷ Number of filters

**for** $j \leftarrow 1$ to $na$ **do**
  $Fimg_j \leftarrow \text{IMFILTER}(Filts_j, img)$ ▷ Set of filtered images
**end for**
**for** $j \leftarrow 1$ to $na/2$ **do**
  $Himg_j \leftarrow Fimg_j + Fimg_{j+na/2}$ ▷ Sum opposite angles
**end for**
$S_+ \leftarrow \sum_1^{na/2} \text{PAIRWMULT}(Himg_j, \text{POSBOOL}(Himg_j))$ ▷ positives sum
$S_- \leftarrow \sum_1^{na/2} \text{PAIRWMULT}(Himg_j, 1 - \text{POSBOOL}(Himg_j))$ ▷ negatives sum
$S_r \leftarrow \text{PAIRWDIV}(S_+, (S_- + 1))$ ▷ Compute ratios
$S_l \leftarrow \text{LOG}(1 - S_r)$ ▷ Take the log
$score \leftarrow \text{EXP}(\text{IMCOMPLEMENT}(S_l))$ ▷ Compute score
$mask \leftarrow \text{PAIRWMULT}(mask0, \text{THRESHOLD}(score, s_0))$ ▷ Final mask
**return** $mask$
**end function**

---

**Algorithm 2:** lowpassfilters function.
Pseudocode for constructing an array of low pass filters. Each filter will compare each value relative to its neighbours, but only in an angle. Constructing the filter using a 2D Gaussian density function makes the filter less sensitive to image noise.

**Require:** Parameters: pixel length $d$, width $w$, and set of angles $A = \{A_1, A_1 + \pi\}$. Some predefined functions: GAUSSPROJ returning the integral of a 2D gaussian density function (with mean $x, y$ and standard deviation $w$) over a space grid; and SUM2 that sums all elements of a matrix.

---

**function** LOWPASSFILTERS($d$,$w$,$A$) ▷ Returns cell array $Filts$
  $na \leftarrow \text{LENGTH}(A)$ ▷ Number of filters
  $ngrid \leftarrow 2d + 1$ ▷ Size of filter
  **for** $j \leftarrow 1$ to $na$ **do** ▷ Compute filter for each angle
    $Filts_j \leftarrow \text{ZEROS}(ngrid, ngrid)$ ▷ Initialise to zeros
    $a \leftarrow A(j)$ ▷ angle
    **for** $q \leftarrow 1$ to $d$ **do** ▷ move from center to $d$
      $x \leftarrow q * \text{COS}(a)$ ▷ X projection
      $x \leftarrow q * \text{SIN}(a)$ ▷ Y projection
      $Filts_j \leftarrow Filts_j + \text{GAUSSPROJ}(x, y, w)$ ▷ Accumulate 2D distributions
    **end for**
    $\theta \leftarrow \text{SUM2}(Filts_j)$ ▷ Sum all values so far
    $Filts_j \leftarrow \theta * \text{GAUSSPROJ}(0, 0, w) - Filts_j$ ▷ Final local difference filter
  **end for**
  **return** $Filts$
**end function**

---

### 3 Mother machine

#### 3.1 Microfluidics device fabrication

The protocol used to fabricate the microfluidics chips is summarised as follows: First, the surface of the master wafer was treated with silane to facilitate removal of PDMS from the surface. The wafer was then taped to the bottom of a large petri dish to secure it in place. Mixed PDMS (1:10 ratio of curing agent to base) was poured onto the master mould to achieve a thickness of approximately 5 mm. The freshly poured PDMS was degassed for 1 h in a vacuum bell jar to remove bubbles followed by curing at 65°C overnight. After cooling to room temperature, chips were then carefully cut out using a scalpel and feeding channels created using a 0.75 mm (ID) biopsy punch (World Precision Instruments Limited). Chips were cleaned by sonicating in isopropanol for 30 min and then left to air dry overnight at 65°C in a closed petri dish (features facing up). Coverslips (Duran, 24x60 mm, #1.5) were cleaned by sonicating in 1 M KOH for 30 min, followed by rinsing three times in Milli-Q water, and then sonication for a further 30 min in Milli-Q water. Coverslips were left to dry overnight at 65°C. Before bonding, chips and coverslips were surface-activated in an oxygen plasma cleaner operated at high intensity under vacuum for 60 s. Bonded chips were then left at 65°C for at least 10 min followed by storage in parafilm-sealed petri dishes at room temperature.

Table 5: **Microchannel dimensions used for different growth media**.

| **Growth Medium** | **Height ($\mu$m)** | **Width ($\mu$m)** | **Length ($\mu$m)** |
|---|---|---|---|
| M9-glucose+amino-acids | 1.36±0.08 | 1.4-1.6 | 26 |
| M9-glucose | 0.91±0.03 | 1.1-1.3 | 25±2 |
| M9-glycerol | 0.91±0.03 | 1.1-1.2 | 25±2 |

#### 3.2 Mother machine data analysis

Additional notes on data curation for mother machine data sets: Initially cells had yet to fully adapt to the imaging conditions as indicated by the mkate2 signal degrading for approximately 2 hours after imaging began, after which it stabilised. This data was discarded and not used for data analysis. At most, 4 hours were truncated from the beginning of data sets. For growth rate and division rate calculations, all non-growing cell cycles (defined by a minimum growth rate of 0.03 $h^{-1}$) were discarded.

### 4 Estimating the transition rate to the high SOS state from mother machine experiments

To analyse the Mother Machine data we used a simple mathematical model (described in the main text). Cells can switch from low to high SOS at rate $\alpha$ and we assume switching back to low SOS is very rare so we neglect this possibility. We assume that this switching can be described by a Poisson process. We estimated $\alpha$ by Maximum likelihood in each nutrient condition using single-cell time-lapse recording of GFP intensity from the Mother Machine. Only mother cells were included in the analysis.

Assuming conversion from low to high SOS can be described by a Poisson process, the time it takes for a given lineage to pass some critical GFP intensity follows an exponential distribution with rate $\alpha$. We call such elapsed time $t_S$. Then, for every lineage beginning at a low-SOS levels (i.e. GFP below the threshold) we distinguish two outcomes: i) the lineage crosses the threshold at a given time from the first observation, ii) the lineage does not cross the threshold over the whole period when it is observed. Given $t_S$ is assumed to follow an exponential distribution, we can estimate the probabilities associated to each event class.

For the first case, we will describe its probability as the probability that $t_S$ falls within the current and previous time interval. Whereas the probability associated to the second case is the probability that $t_S$ is larger than the total observed time for that lineage (i.e. 1 minus the c.d.f of $t_S$). Then

| **Case** | **Description** | **Probability** |
|---|---|---|
| i | Lineage crosses GFP theshold at time $t$ | $\mathbb{P}(t - t_\Delta > t_S \geq t \vert \alpha) = e^{-\alpha t}\left(e^{\alpha t_\Delta} - 1\right)$ |
| ii | Lineage does not crosses GFP theshold at time $t$ | $\mathbb{P}(t_S > t \vert \alpha) = 1 - (1 - e^{-\alpha t}) = e^{-\alpha t}$ |

where $t_\Delta$ is the time interval used for imaging.

With these definitions, we classify each lineage into both cases, and define a likelihood for each set of observations. Let's call the set of all elapsed times for the lineages in the first case by $T_1$, and the total observed time for all lineages in the second case by $T_2$. Then we define the likelihood of our observation as the product of the probabilities of their respective cases

$$\mathcal{L} = \mathcal{L}_1\mathcal{L}_2 = \prod_{t_1 \in T_1} \mathbb{P}(t_1 - t_\Delta > t_S \geq t_2|\alpha) \prod_{t_2 \in T_2} \mathbb{P}(t_S > t_2|\alpha)$$

where the log-likelihood is

$$\ln \mathcal{L}(\alpha) = |T_1| \ln\left(e^{\alpha t_\Delta} - 1\right) - \alpha \left(\sum_{t_1 \in T_1} t_1 + \sum_{t_2 \in T_2} t_2\right) \tag{1}$$

where $|T_1|$ is the size of the set $T_1$, that is all observed cases that do cross the threshold. Also, using the partial derivative the log-likelihood we can obtain the value of $\alpha$ with maximum likelihood ($\alpha_m$), that is

$$\alpha_m = \frac{1}{t_\Delta} \ln\left(\frac{\sum_{t_1 \in T_1} t_1 + \sum_{t_2 \in T_2} t_2}{\sum_{t_1 \in T_1} t_1 + \sum_{t_2 \in T_2} t_2 - |T_1| t_\Delta}\right) \tag{2}$$

In order to estimate the confidence in the estimation of $\alpha$, we uzed the Metropolis-Hastings algorithm starting form $\alpha_m$. During each iteration of the Markov chain, candidates were generated by sampling $\alpha' = \alpha_{i-1} + \delta$ where the index $i$ represent the previous iteration, and

$\delta \sim uniform([-\epsilon, \epsilon])$. Whenever some sampled candidate $\alpha'$ was found to be negative, $\alpha'$ was sampled again until it was not longer so. The acceptance ratio was computed using the log-likelihood, and was equal to $\exp(\ln \mathcal{L}(\alpha') - \ln \mathcal{L}(\alpha_i))$. Monte-Carlo simulations were performed through $10^5$ iterations using $\epsilon = 10^{-3}$.

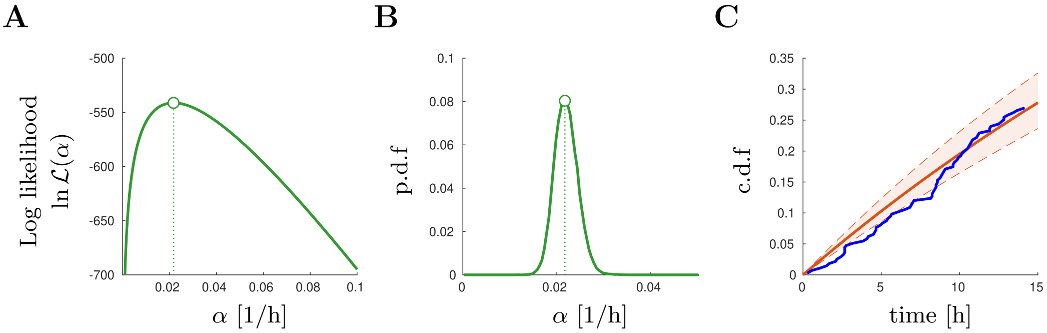


Figure 2: **Estimation of the switching rate $\alpha$ from mother-machine experiments**. Given a GFP intensity (SOS) threshold, we assume that the conversion between low and high SOS can be described by a Poisson process which allows us to derive explicit expressions for the likelihood of SOS transitions based on the transition rate $\alpha$. In particular, we can obtain the value of $\alpha$ with maximum likelihood (panel A), and use Metropolis-Hastings algorithm to estimate the p.d.f of $\alpha$ (panel B). In panel C we compare the empirical cumulative density function of elapsed times to pass the given threshold (blue line), with the expected exponential distribution (red line). The green shared area represents the 90% confidence interval based on the p.d.f estimates of $\alpha$. This example was constructed using a GFP threshold of 8 A.U. using one of the dataset for the strain carrying 2 palindromes in glucose + amino-acids condition.

### 5 Estimating the steady-state population growth-rate from single-cell division time distributions

Different division times distributions will result in populations "effectively" growing at different rates, and to compute this value taking the average division time is not accurate (Painter and Marr, 1968; Thomas, 2017). To avoid confusions, we wish to clarify that we are referring here specifically to the growth-rate of the population size in number of cells.

The relation between the distribution of division times and the population growth-rate is given by a functional equation and cannot be calculated explicitly (Painter and Marr, 1968; Thomas, 2017). Let's call $\tau$ a random variable denoting the division time which follows a $\phi(\tau)$ distribution and $\lambda$ the steady-state population growth-rate. The population growth-rate $\lambda$ satisfies the following equation assuming that correlation between successive division event can be neglected:

$$1 = 2 \int_0^\infty \phi(\tau)\ e^{-\lambda \tau} d\tau \tag{3}$$

In order to estimate $\lambda$ from our experimental measurements, we follow a least-square strategy where $\lambda$ is the value that minimizes the distance to the expected relation in equation 3. However, before we do this we need to estimate the distribution of division times $\phi(\tau)$ from our experimental measurements, taking into account that our measurements of division times are discrete due to the imaging protocol.

To simplify the problem, we divide $\phi(\tau)$ into separate intervals with width identical to the imaging acquisition interval (called $t_\Delta$, and assume that the probability of observing a division event within that interval is uniform. We will use here an index $j \in \mathbb{N}_0^+$ to denote each interval. Finally, we make the simplification that the observed frequency of division events approximates the actual distribution. Then, combining these assumptions we have

$$\phi(\tau) \approx \frac{f(j)}{t_\Delta},$$

where $f(j)$ is the relative frequency of division events within an interval $j$. Note that for a given value of $\tau$ the corresponding interval will be different: $(j-1)t_\Delta < \tau \leq jt_\Delta$. Now we can combine our simplifications to replace the right side term of equation 3 and obtain

$$\int_0^\infty \phi(\tau)\, e^{-\lambda\tau} d\tau \; \approx \sum_{j=1}^{\infty} \frac{f(j)}{t_\Delta} \int_{(j-1)t_\Delta}^{jt_\Delta} e^{-\lambda\tau} d\tau = \frac{1}{\lambda t_\Delta}\left(e^{\lambda t_\Delta} - 1\right) \sum_{j=1}^{\infty} f(j) e^{-j\lambda t_\Delta}$$

Then, we estimate $\lambda$ as the value that minimizes the following expression:

$$\lambda = \underset{\lambda \in \mathbb{R}^+}{\text{minimize}} \left(1 - 2\frac{1}{\lambda t_\Delta}\left(e^{\lambda t_\Delta} - 1\right) \sum_{j=1}^{\infty} f(j) e^{-j\lambda t_\Delta}\right)^2$$

Figure 3 shows that the minimal $\lambda$ value is unique, and also reproducible between experimental repeats.

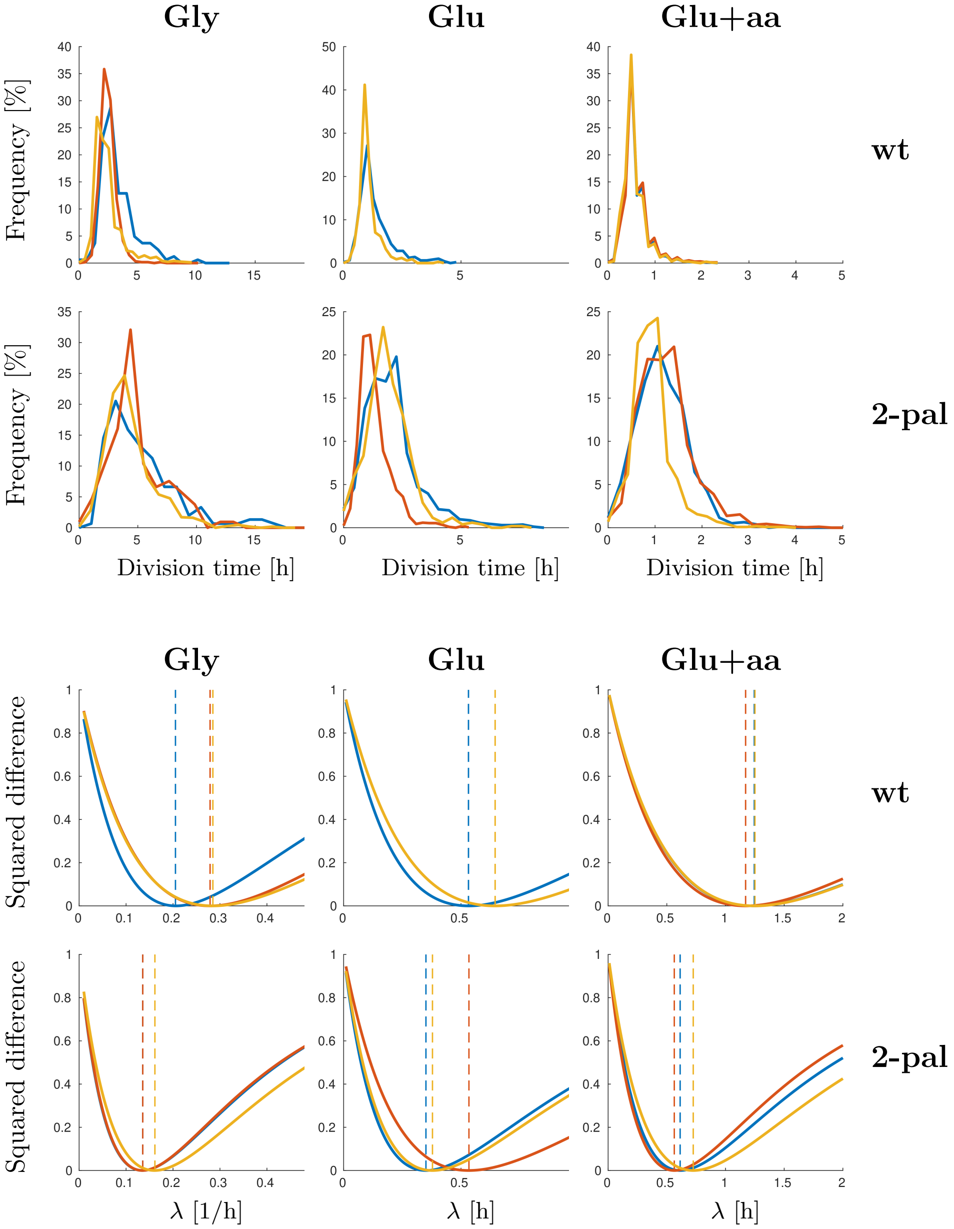


Figure 3: **Estimation of population growth-rate from single-cell divisions**. On the top, we display the division-time distributions for wild-type and 2-pal strains in three different media. On the bottom, we show the cost function associated to a given growth-rate value. The growth-rate with minimum value in the curve (dashed-line) corresponds to the estimated population growth-rate given the division time statistics. Each color represents one biological repeat.

#### 6 Estimating the fraction of high SOS cell in a growing population

Let's consider $n_1$ and $n_2$ the number of cells in each state, and $\lambda_1$ and $\lambda_2$ the respective division rates. We call $n_1$ the number of "healthy" low SOS cells, and $n_2$ the number of slow-dividing cells with high SOS induction. Let us assume first-order kinetics for conversion between the two populations, and call $\alpha$ the conversion rate constant from population one to population two, and $\beta$ the rate constant for the reverse reaction. Thus, the population dynamics is given by the following equations

$$\begin{aligned}\frac{dn_1}{dt} &= (\lambda_1-\alpha)n_1+\beta n_2\\ \frac{dn_2}{dt} &= (\lambda_2-\beta)n_2+\alpha n_1\end{aligned}$$

We are interested in the population fractions and call them $f_1=\frac{n_1}{n_1+n_2}\in[0,1]$, and $f_2=\frac{n_2}{n_1+n_2}\in[0,1]$. Converting the dynamics into fractions we get:

$$\begin{aligned}\frac{df_1}{dt} &= -\alpha f_1+\beta f_2+f_1f_2(\lambda_1-\lambda_2)\\ \frac{df_2}{dt} &= \alpha f_1-\beta f_2-f_1f_2(\lambda_1-\lambda_2)\end{aligned} \tag{4}$$

Notice that the total population growth is given by:

$$\frac{d(n_1+n_2)}{dt}=(\lambda_1f_1+\lambda_2f_2)(n_1+n_2) \tag{5}$$

At steady-state, the population fractions are time-invariant. The solution for $f_1$ and $f_2$ at steady state is given by:

$$\begin{aligned}f_1 &= \frac{1}{2}\left(1+\frac{\alpha+\beta\pm\sqrt{\delta}}{\lambda_2-\lambda_1}\right)\\ f_2 &= \frac{1}{2}\left(1-\frac{\alpha+\beta\pm\sqrt{\delta}}{\lambda_2-\lambda_1}\right)\end{aligned} \tag{6}$$

where $\delta=(\beta+\alpha+\lambda_2-\lambda_1)^2-4\beta(\lambda_2-\lambda_1)$. The steady-state can be approximated when $\alpha\gg\beta$ (and therefore $\beta$ is negligible as we assumed previously) and $\lambda_1\gg\lambda_2$. Under this hypotheses we have:

$$\begin{aligned}f_2 &\approx \alpha/\lambda_1\\ f_1 &\approx 1-\alpha/\lambda_1\end{aligned} \tag{7}$$

Therefore, using this model, we can predict $f_2$, the fraction of high SOS cells based on the parameters measured in the Mother Machine and compare this prediction to the experimentally measured fraction of high SOS cells (see main text).
