## Supplementary data: Agar-pad snapshot values for "Growth-dependent heterogeneity in the DNA damage response in *Escherichia coli*"

List of values as support in the main text

| Metric | strainID | Strain | Cipro | media avg gly | avg glu | avg glu+aa | err gly | err glu | err glu+aa |
|---|---|---|---|---|---|---|---|---|---|
| doubling rate [dlb/hr] | eSJR206 | wt | 0 | 0,6 | 1,04 | 1,61 | 0,01 | 0,04 | 0,05 |
| doubling rate [dlb/hr] | eSJR301 | r-pal | 0 | 0,5 | 0,89 | 1,5 | 0,01 | 0,03 | 0,03 |
| doubling rate [dlb/hr] | eSJR214 | l-pal | 0 | 0,59 | 0,96 | 1,54 | 0,02 | 0,05 | 0,04 |
| doubling rate [dlb/hr] | eSJR302 | 2-pal | 0 | 0,45 | 0,81 | 1,42 | 0,02 | 0,02 | 0,03 |
| doubling rate [dlb/hr] | eSJR206 | wt | 1 | 0,58 | 1,01 | 1,6 | 0,01 | 0,03 | 0,04 |
| doubling rate [dlb/hr] | eSJR206 | wt | 2 | 0,57 | 0,93 | 1,61 | 0,01 | 0,03 | 0,04 |
| doubling rate [dlb/hr] | eSJR206 | wt | 3 | 0,553 | 0,91 | 1,54 | 0,01 | 0,03 | 0,04 |
| | | | | **Number of repeats** | | | **Number of cells per repeat** | | |
| Repeats / sample size | eSJR203 | wt-auto | 0 | 4 | 3 | 3 | 39778 116 | 32196 138 | 7901 15398 9246 |
| Repeats / sample size | eSJR292 | lexA3 | 0 | 3 | 3 | 3 | 21784 693 | 15484 130 | 6401 7702 8015 |
| Repeats / sample size | eSJR206 | wt | 0 | 6 | 9 | 7 | 32201 715 | 23205 135 | 12725 7333 16817 99 |
| Repeats / sample size | eSJR301 | r-pal | 0 | 3 | 3 | 3 | 13768 130 | 20502 137 | 14477 14802 7328 |
| Repeats / sample size | eSJR214 | l-pal | 0 | 3 | 3 | 3 | 28141 546 | 19715 133 | 6101 6514 9892 |
| Repeats / sample size | eSJR302 | 2-pal | 0 | 3 | 3 | 4 | 30226 295 | 17255 195 | 7803 4638 8822 9806 |
| Repeats / sample size | eSJR206 | wt | 1 | 3 | 3 | 4 | 25231 735 | 12755 994 | 7472 8708 14488 804 |
| Repeats / sample size | eSJR206 | wt | 2 | 3 | 3 | 4 | 17157 514 | 5927 1162 | 5279 8880 7517 5349 |
| Repeats / sample size | eSJR206 | wt | 3 | 3 | 3 | 4 | 7924 1508 | 5733 7158 | 8193 2097 5706 3566 |
| | | | | **PsfiA-mGFP mean and mode** | | | | | |
| Mean PsfiA-GFP [a.u.] | eSJR203 | wt-auto | 0 | 0,79 | 0,83 | 0,82 | 0,02 | 0,02 | 0,01 |
| Mean PsfiA-GFP [a.u.] | eSJR292 | lexA3 | 0 | 0,85 | 0,89 | 0,96 | 0,01 | 0,02 | 0,05 |
| Mean PsfiA-GFP [a.u.] | eSJR206 | wt | 0 | 0,98 | 0,98 | 1,04 | 0,02 | 0,01 | 0,05 |
| Mean PsfiA-GFP [a.u.] | eSJR301 | r-pal | 0 | 1,53 | 1,61 | 1,6 | 0,07 | 0,04 | 0,1 |
| Mean PsfiA-GFP [a.u.] | eSJR214 | l-pal | 0 | 2,5 | 2,47 | 1,9 | 0,08 | 0,07 | 0,1 |
| Mean PsfiA-GFP [a.u.] | eSJR302 | 2-pal | 0 | 3,1 | 2,9 | 2,5 | 0,2 | 0,2 | 0,1 |
| Mean PsfiA-GFP [a.u.] | eSJR206 | wt | 1 | 1,6 | 1,73 | 1,39 | 0,1 | 0,07 | 0,07 |
| Mean PsfiA-GFP [a.u.] | eSJR206 | wt | 2 | 2,6 | 2,8 | 1,84 | 0,1 | 0,1 | 0,09 |
| Mean PsfiA-GFP [a.u.] | eSJR206 | wt | 3 | 3,8 | 3,6 | 2,37 | 0,3 | 0,1 | 0,1 |
| Mode PsfiA-GFP [a.u.] | eSJR203 | wt-auto | 0 | 0,86 | 0,91 | 0,9 | 0,02 | 0,02 | 0,02 |
| Mode PsfiA-GFP [a.u.] | eSJR292 | lexA3 | 0 | 0,96 | 1,01 | 1,08 | 0,01 | 0,02 | 0,06 |
| Mode PsfiA-GFP [a.u.] | eSJR206 | wt | 0 | 1,06 | 1,08 | 1,2 | 0,02 | 0,01 | 0,08 |
| Mode PsfiA-GFP [a.u.] | eSJR301 | r-pal | 0 | 1,31 | 1,58 | 1,9 | 0,02 | 0,02 | 0,1 |
| Mode PsfiA-GFP [a.u.] | eSJR214 | l-pal | 0 | 1,55 | 1,76 | 2 | 0,07 | 0,07 | 0,1 |
| Mode PsfiA-GFP [a.u.] | eSJR302 | 2-pal | 0 | 1,8 | 2,2 | 2,7 | 0,1 | 0,2 | 0,1 |
| Mode PsfiA-GFP [a.u.] | eSJR206 | wt | 1 | 1,7 | 1,84 | 1,62 | 0,1 | 0,07 | 0,08 |
| Mode PsfiA-GFP [a.u.] | eSJR206 | wt | 2 | 2,3 | 2,81 | 2,15 | 0,1 | 0,06 | 0,1 |
| Mode PsfiA-GFP [a.u.] | eSJR206 | wt | 3 | 3,3 | 3,83 | 2,68 | 0,3 | 0,09 | 0,07 |
| | | | | **Average PsfiA-mGFP above/bellow population percentile [a.u.]** | | | | | |
| Bottom 99 PsfiA-GFP [a.u.] | eSJR203 | wt-auto | 0 | 0,79 | 0,83 | 0,82 | 0,02 | 0,02 | 0,01 |
| Bottom 99 PsfiA-GFP [a.u.] | eSJR292 | lexA3 | 0 | 0,85 | 0,88 | 0,95 | 0,01 | 0,02 | 0,05 |
| Bottom 99 PsfiA-GFP [a.u.] | eSJR206 | wt | 0 | 0,93 | 0,95 | 1,02 | 0,02 | 0,01 | 0,05 |
| Bottom 85 PsfiA-GFP [a.u.] | eSJR301 | r-pal | 0 | 1,07 | 1,23 | 1,4 | 0,03 | 0,02 | 0,1 |
| Bottom 85 PsfiA-GFP [a.u.] | eSJR214 | l-pal | 0 | 1,34 | 1,44 | 1,53 | 0,07 | 0,06 | 0,1 |
| Bottom 85 PsfiA-GFP [a.u.] | eSJR302 | 2-pal | 0 | 1,7 | 1,8 | 2,01 | 0,2 | 0,1 | 0,08 |
| Bottom 85 PsfiA-GFP [a.u.] | eSJR206 | wt | 1 | 1,26 | 1,36 | 1,23 | 0,09 | 0,04 | 0,06 |
| Bottom 85 PsfiA-GFP [a.u.] | eSJR206 | wt | 2 | 1,74 | 2,09 | 1,59 | 0,09 | 0,04 | 0,08 |
| Bottom 85 PsfiA-GFP [a.u.] | eSJR206 | wt | 3 | 2,5 | 2,81 | 2 | 0,2 | 0,07 | 0,06 |
| Top 1 PsfiA-GFP [a.u.] | eSJR203 | wt-auto | 0 | 0,9 | 0,95 | 0,95 | 0,02 | 0,03 | 0,02 |
| Top 1 PsfiA-GFP [a.u.] | eSJR292 | lexA3 | 0 | 1,3 | 1,14 | 1,4 | 0,05 | 0,03 | 0,3 |
| Top 1 PsfiA-GFP [a.u.] | eSJR206 | wt | 0 | 5,6 | 3,9 | 2,7 | 0,4 | 0,2 | 0,2 |
| Top 15 PsfiA-GFP [a.u.] | eSJR301 | r-pal | 0 | 4,2 | 3,7 | 3 | 0,3 | 0,1 | 0,3 |
| Top 15 PsfiA-GFP [a.u.] | eSJR214 | l-pal | 0 | 9,1 | 8,3 | 4 | 0,3 | 0,2 | 0,3 |
| Top 15 PsfiA-GFP [a.u.] | eSJR302 | 2-pal | 0 | 11 | 9,5 | 5,3 | 0,5 | 0,7 | 0,4 |
| Top 15 PsfiA-GFP [a.u.] | eSJR206 | wt | 1 | 3,7 | 3,8 | 2,3 | 0,2 | 0,2 | 0,1 |
| Top 15 PsfiA-GFP [a.u.] | eSJR206 | wt | 2 | 7,2 | 6,5 | 3,2 | 0,5 | 0,5 | 0,2 |
| Top 15 PsfiA-GFP [a.u.] | eSJR206 | wt | 3 | 10,9 | 8,1 | 4,5 | 0,7 | 0,5 | 0,4 |
| | | | | **Population fractions above Psfi-GFP threshold [%]** | | | | | |
| Pop fraction PsfiA-GFP >= 2 [a.u.] | eSJR203 | wt-auto | 0 | 0 | 0,002 | 0 | 0 | 0,002 | 0 |
| Pop fraction PsfiA-GFP >= 2 [a.u.] | eSJR292 | lexA3 | 0 | 0,078 | 0 | 0,06 | 0,009 | 0 | 0,04 |
| Pop fraction PsfiA-GFP >= 2 [a.u.] | eSJR206 | wt | 0 | 1,2 | 0,94 | 1,1 | 0,1 | 0,06 | 0,4 |
| Pop fraction PsfiA-GFP >= 2 [a.u.] | eSJR301 | r-pal | 0 | 10,1 | 13 | 19 | 0,9 | 1 | 6 |
| Pop fraction PsfiA-GFP >= 2 [a.u.] | eSJR214 | l-pal | 0 | 23 | 26 | 26 | 3 | 2 | 6 |
| Pop fraction PsfiA-GFP >= 2 [a.u.] | eSJR302 | 2-pal | 0 | 36 | 40 | 55 | 6 | 6 | 4 |
| Pop fraction PsfiA-GFP >= 2 [a.u.] | eSJR206 | wt | 1 | 12 | 19 | 7 | 3 | 3 | 2 |
| Pop fraction PsfiA-GFP >= 2 [a.u.] | eSJR206 | wt | 2 | 38 | 61 | 27 | 5 | 2 | 5 |
| Pop fraction PsfiA-GFP >= 2 [a.u.] | eSJR206 | wt | 3 | 74 | 90 | 56 | 8 | 0,7 | 4 |
| Pop fraction PsfiA-GFP >= 3 [a.u.] | eSJR203 | wt-auto | 0 | 0 | 0 | 0 | 0 | 0 | 0 |
| Pop fraction PsfiA-GFP >= 3 [a.u.] | eSJR292 | lexA3 | 0 | 0,04 | 0 | 0,05 | 0,01 | 0 | 0,03 |
| Pop fraction PsfiA-GFP >= 3 [a.u.] | eSJR206 | wt | 0 | 0,6 | 0,31 | 0,2 | 0,07 | 0,02 | 0,06 |
| Pop fraction PsfiA-GFP >= 3 [a.u.] | eSJR301 | r-pal | 0 | 6,2 | 4,9 | 4 | 0,4 | 0,5 | 2 |
| Pop fraction PsfiA-GFP >= 3 [a.u.] | eSJR214 | l-pal | 0 | 14,7 | 13,2 | 8 | 0,8 | 0,9 | 2 |
| Pop fraction PsfiA-GFP >= 3 [a.u.] | eSJR302 | 2-pal | 0 | 23 | 19 | 19 | 3 | 3 | 3 |
| Pop fraction PsfiA-GFP >= 3 [a.u.] | eSJR206 | wt | 1 | 3,8 | 5 | 1,4 | 0,5 | 0,9 | 0,3 |
| Pop fraction PsfiA-GFP >= 3 [a.u.] | eSJR206 | wt | 2 | 13 | 20 | 4,9 | 1 | 2 | 0,8 |
| Pop fraction PsfiA-GFP >= 3 [a.u.] | eSJR206 | wt | 3 | 36 | 47 | 12 | 8 | 3 | 2 |
| Pop fraction PsfiA-GFP >= 5 [a.u.] | eSJR203 | wt-auto | 0 | 0 | 0 | 0 | 0 | 0 | 0 |
| Pop fraction PsfiA-GFP >= 5 [a.u.] | eSJR292 | lexA3 | 0 | 0,014 | 0 | 0,02 | 0,008 | 0 | 0,02 |
| Pop fraction PsfiA-GFP >= 5 [a.u.] | eSJR206 | wt | 0 | 0,3 | 0,14 | 0,05 | 0,03 | 0,01 | 0,01 |
| Pop fraction PsfiA-GFP >= 5 [a.u.] | eSJR301 | r-pal | 0 | 4,1 | 2,6 | 1 | 0,3 | 0,2 | 0,2 |
| Pop fraction PsfiA-GFP >= 5 [a.u.] | eSJR214 | l-pal | 0 | 9,8 | 7,3 | 2,3 | 0,4 | 0,2 | 0,4 |
| Pop fraction PsfiA-GFP >= 5 [a.u.] | eSJR302 | 2-pal | 0 | 14 | 9 | 4,3 | 1 | 1 | 0,7 |
| Pop fraction PsfiA-GFP >= 5 [a.u.] | eSJR206 | wt | 1 | 1,9 | 1,7 | 0,47 | 0,1 | 0,2 | 0,03 |
| Pop fraction PsfiA-GFP >= 5 [a.u.] | eSJR206 | wt | 2 | 6,1 | 5,5 | 1,1 | 0,4 | 0,7 | 0,2 |
| Pop fraction PsfiA-GFP >= 5 [a.u.] | eSJR206 | wt | 3 | 13 | 9 | 2,4 | 2 | 1 | 0,3 |
| Pop fraction PsfiA-GFP >= 7.5 [a.u.] | eSJR203 | wt-auto | 0 | 0 | 0 | 0 | 0 | 0 | 0 |
| Pop fraction PsfiA-GFP >= 7.5 [a.u.] | eSJR292 | lexA3 | 0 | 0,005 | 0 | 0,02 | 0,005 | 0 | 0,02 |
| Pop fraction PsfiA-GFP >= 7.5 [a.u.] | eSJR206 | wt | 0 | 0,18 | 0,1 | 0,024 | 0,03 | 0,01 | 0,008 |
| Pop fraction PsfiA-GFP >= 7.5 [a.u.] | eSJR301 | r-pal | 0 | 2,4 | 1,6 | 0,37 | 0,2 | 0,07 | 0,08 |
| Pop fraction PsfiA-GFP >= 7.5 [a.u.] | eSJR214 | l-pal | 0 | 6,9 | 5,1 | 1,1 | 0,5 | 0,1 | 0,1 |
| Pop fraction PsfiA-GFP >= 7.5 [a.u.] | eSJR302 | 2-pal | 0 | 9,4 | 6,3 | 1,8 | 0,6 | 0,6 | 0,2 |
| Pop fraction PsfiA-GFP >= 7.5 [a.u.] | eSJR206 | wt | 1 | 1,28 | 1,1 | 0,2 | 0,06 | 0,1 | 0,03 |
| Pop fraction PsfiA-GFP >= 7.5 [a.u.] | eSJR206 | wt | 2 | 4,1 | 3,1 | 0,49 | 0,4 | 0,4 | 0,09 |
| Pop fraction PsfiA-GFP >= 7.5 [a.u.] | eSJR206 | wt | 3 | 7,8 | 4,3 | 1,2 | 0,6 | 0,5 | 0,2 |
| Pop fraction PsfiA-GFP >= 10 [a.u.] | eSJR203 | wt-auto | 0 | 0 | 0 | 0 | 0 | 0 | 0 |
| Pop fraction PsfiA-GFP >= 10 [a.u.] | eSJR292 | lexA3 | 0 | 0 | 0 | 0 | 0 | 0 | 0 |
| Pop fraction PsfiA-GFP >= 10 [a.u.] | eSJR206 | wt | 0 | 0,13 | 0,08 | 0,014 | 0,02 | 0,01 | 0,004 |
| Pop fraction PsfiA-GFP >= 10 [a.u.] | eSJR301 | r-pal | 0 | 1,2 | 0,91 | 0,13 | 0,2 | 0,02 | 0,04 |
| Pop fraction PsfiA-GFP >= 10 [a.u.] | eSJR214 | l-pal | 0 | 4,8 | 3,9 | 0,67 | 0,4 | 0,1 | 0,07 |
| Pop fraction PsfiA-GFP >= 10 [a.u.] | eSJR302 | 2-pal | 0 | 6,5 | 4,6 | 1,1 | 0,4 | 0,4 | 0,2 |
| Pop fraction PsfiA-GFP >= 10 [a.u.] | eSJR206 | wt | 1 | 0,93 | 0,73 | 0,12 | 0,05 | 0,07 | 0,02 |
| Pop fraction PsfiA-GFP >= 10 [a.u.] | eSJR206 | wt | 2 | 3 | 2,1 | 0,26 | 0,3 | 0,3 | 0,06 |
| Pop fraction PsfiA-GFP >= 10 [a.u.] | eSJR206 | wt | 3 | 5,6 | 2,9 | 0,8 | 0,5 | 0,3 | 0,2 |
| Pop fraction PsfiA-GFP >= 12.5 [a.u.] | eSJR203 | wt-auto | 0 | 0 | 0 | 0 | 0 | 0 | 0 |
| Pop fraction PsfiA-GFP >= 12.5 [a.u.] | eSJR292 | lexA3 | 0 | 0 | 0 | 0 | 0 | 0 | 0 |
| Pop fraction PsfiA-GFP >= 12.5 [a.u.] | eSJR206 | wt | 0 | 0,09 | 0,053 | 0,01 | 0,02 | 0,008 | 0,004 |
| Pop fraction PsfiA-GFP >= 12.5 [a.u.] | eSJR301 | r-pal | 0 | 0,5 | 0,45 | 0,04 | 0,1 | 0,01 | 0,02 |
| Pop fraction PsfiA-GFP >= 12.5 [a.u.] | eSJR214 | l-pal | 0 | 3,4 | 2,95 | 0,45 | 0,2 | 0,09 | 0,07 |
| Pop fraction PsfiA-GFP >= 12.5 [a.u.] | eSJR302 | 2-pal | 0 | 4,4 | 3,5 | 0,7 | 0,4 | 0,3 | 0,1 |
| Pop fraction PsfiA-GFP >= 12.5 [a.u.] | eSJR206 | wt | 1 | 0,69 | 0,53 | 0,07 | 0,04 | 0,05 | 0,01 |
| Pop fraction PsfiA-GFP >= 12.5 [a.u.] | eSJR206 | wt | 2 | 2,1 | 1,5 | 0,14 | 0,2 | 0,3 | 0,02 |
| Pop fraction PsfiA-GFP >= 12.5 [a.u.] | eSJR206 | wt | 3 | 4,1 | 2,2 | 0,6 | 0,3 | 0,2 | 0,2 |
| Pop fraction PsfiA-GFP >= 15 [a.u.] | eSJR203 | wt-auto | 0 | 0 | 0 | 0 | 0 | 0 | 0 |

| | | | | | | | | | |
|---|---|---|---|---|---|---|---|---|---|
| Pop fraction PsfiA-GFP >= 15 [a.u.] | eSJR292 | lexA3 | 0 | 0 | 0 | 0 | 0 | 0 | 0 |
| Pop fraction PsfiA-GFP >= 15 [a.u.] | eSJR206 | wt | 0 | 0,07 | 0,037 | 0,006 | 0,01 | 0,005 | 0,002 |
| Pop fraction PsfiA-GFP >= 15 [a.u.] | eSJR301 | r-pal | 0 | 0,26 | 0,222 | 0,014 | 0,05 | 0,009 | 0,008 |
| Pop fraction PsfiA-GFP >= 15 [a.u.] | eSJR214 | l-pal | 0 | 2,4 | 2,29 | 0,3 | 0,2 | 0,08 | 0,06 |
| Pop fraction PsfiA-GFP >= 15 [a.u.] | eSJR302 | 2-pal | 0 | 3,1 | 2,7 | 0,5 | 0,3 | 0,3 | 0,1 |
| Pop fraction PsfiA-GFP >= 15 [a.u.] | eSJR206 | wt | 1 | 0,49 | 0,39 | 0,039 | 0,06 | 0,05 | 0,006 |
| Pop fraction PsfiA-GFP >= 15 [a.u.] | eSJR206 | wt | 2 | 1,5 | 1,1 | 0,07 | 0,2 | 0,3 | 0,02 |
| Pop fraction PsfiA-GFP >= 15 [a.u.] | eSJR206 | wt | 3 | 3 | 1,6 | 0,4 | 0,3 | 0,2 | 0,2 |

**Constitutive expression [A.U.]**

| | | | | | | | | | |
|---|---|---|---|---|---|---|---|---|---|
| Mean PtetO-mKate [a.u.] | eSJR203 | wt-auto | 0 | 1,89 | 1,56 | 0,96 | 0,1 | 0,05 | 0,02 |
| Mean PtetO-mKate [a.u.] | eSJR292 | lexA3 | 0 | 1,89 | 1,47 | 1,01 | 0,05 | 0,04 | 0,02 |
| Mean PtetO-mKate [a.u.] | eSJR206 | wt | 0 | 1,8 | 1,47 | 1 | 0,04 | 0,03 | 0,06 |
| Mean PtetO-mKate [a.u.] | eSJR301 | r-pal | 0 | 2,11 | 1,62 | 1,18 | 0,04 | 0,02 | 0,05 |
| Mean PtetO-mKate [a.u.] | eSJR214 | l-pal | 0 | 1,94 | 1,58 | 1,05 | 0,07 | 0,03 | 0,04 |
| Mean PtetO-mKate [a.u.] | eSJR302 | 2-pal | 0 | 2,6 | 1,83 | 1,32 | 0,1 | 0,01 | 0,05 |
| Mean PtetO-mKate [a.u.] | eSJR206 | wt | 1 | 1,9 | 1,54 | 1,01 | 0,2 | 0,03 | 0,06 |
| Mean PtetO-mKate [a.u.] | eSJR206 | wt | 2 | 1,9 | 1,54 | 0,96 | 0,1 | 0,06 | 0,07 |
| Mean PtetO-mKate [a.u.] | eSJR206 | wt | 3 | 2,1 | 1,58 | 1,06 | 0,3 | 0,02 | 0,05 |

**Cell size**

| | | | | | | | | | |
|---|---|---|---|---|---|---|---|---|---|
| Mean cell size [um^2] | eSJR203 | wt-auto | 0 | 1,56 | 1,71 | 2,67 | 0,03 | 0,02 | 0,07 |
| Mean cell size [um^2] | eSJR292 | lexA3 | 0 | 1,46 | 1,69 | 2,5 | 0,02 | 0,05 | 0,2 |
| Mean cell size [um^2] | eSJR206 | wt | 0 | 1,49 | 1,68 | 2,47 | 0,04 | 0,02 | 0,1 |
| Mean cell size [um^2] | eSJR301 | r-pal | 0 | 1,62 | 1,73 | 2,61 | 0,04 | 0,01 | 0,07 |
| Mean cell size [um^2] | eSJR214 | l-pal | 0 | 1,79 | 1,98 | 3,2 | 0,09 | 0,01 | 0,07 |
| Mean cell size [um^2] | eSJR302 | 2-pal | 0 | 1,68 | 1,99 | 3,78 | 0,08 | 0,05 | 0,08 |
| Mean cell size [um^2] | eSJR206 | wt | 1 | 1,66 | 2,03 | 3,2 | 0,02 | 0,06 | 0,1 |
| Mean cell size [um^2] | eSJR206 | wt | 2 | 1,94 | 2,83 | 4,3 | 0,04 | 0,09 | 0,1 |
| Mean cell size [um^2] | eSJR206 | wt | 3 | 2,3 | 4,67 | 6,4 | 0,1 | 0,07 | 0,4 |

**Estimation of alpha for different SOS thesholds**

| | strainID | Strain | Cipro | alpha [1/h] | err |
|---|---|---|---|---|---|
| Model PsfiA-GFP threshold 4 | eSJR206 | wt | 0 | 0,0015 | 0,0002 |
| Model PsfiA-GFP threshold 4 | eSJR301 | r-pal | 0 | 0,021 | 0,003 |
| Model PsfiA-GFP threshold 4 | eSJR214 | l-pal | 0 | 0,052 | 0,008 |
| Model PsfiA-GFP threshold 4 | eSJR302 | 2-pal | 0 | 0,08 | 0,01 |
| Model PsfiA-GFP threshold 4 | eSJR206 | wt | 1 | 0,011 | 0,003 |
| Model PsfiA-GFP threshold 4 | eSJR206 | wt | 2 | 0,04 | 0,01 |
| Model PsfiA-GFP threshold 4 | eSJR206 | wt | 3 | 0,09 | 0,03 |

| | | | | Relative distance to model [%] | | | error | | |
|---|---|---|---|---|---|---|---|---|---|
| Model PsfiA-GFP threshold 4 | eSJR206 | wt | 0 | gly | glu | glu+aa | gly | glu | glu+aa |
| Model PsfiA-GFP threshold 4 | eSJR301 | r-pal | 0 | 10 | -14 | -40 | 10 | 7 | 10 |
| Model PsfiA-GFP threshold 4 | eSJR214 | l-pal | 0 | -2 | 9 | -10 | 7 | 8 | 30 |
| Model PsfiA-GFP threshold 4 | eSJR302 | 2-pal | 0 | -6 | 27 | -20 | 3 | 6 | 20 |
| Model PsfiA-GFP threshold 4 | eSJR206 | wt | 1 | -10 | 10 | 10 | 10 | 20 | 20 |
| Model PsfiA-GFP threshold 4 | eSJR206 | wt | 2 | -12 | 60 | -37 | 8 | 20 | 5 |
| Model PsfiA-GFP threshold 4 | eSJR206 | wt | 3 | -14 | 70 | -44 | 6 | 20 | 7 |
| | | | | -10 | 50 | -47 | 20 | 20 | 5 |

| | | | | alpha | err |
|---|---|---|---|---|---|
| Model PsfiA-GFP threshold 6 | eSJR206 | wt | 0 | 0,0009 | 0,0001 |
| Model PsfiA-GFP threshold 6 | eSJR301 | r-pal | 0 | 0,014 | 0,002 |
| Model PsfiA-GFP threshold 6 | eSJR214 | l-pal | 0 | 0,035 | 0,006 |
| Model PsfiA-GFP threshold 6 | eSJR302 | 2-pal | 0 | 0,048 | 0,007 |
| Model PsfiA-GFP threshold 6 | eSJR206 | wt | 1 | 0,007 | 0,002 |
| Model PsfiA-GFP threshold 6 | eSJR206 | wt | 2 | 0,021 | 0,005 |
| Model PsfiA-GFP threshold 6 | eSJR206 | wt | 3 | 0,039 | 0,008 |

| | | | | Relative distance to model [%] | | | error | | |
|---|---|---|---|---|---|---|---|---|---|
| | | | | gly | glu | glu+aa | gly | glu | glu+aa |
| Model PsfiA-GFP threshold 6 | eSJR206 | wt | 0 | 10 | -7 | -60 | 8 | 9 | 10 |
| Model PsfiA-GFP threshold 6 | eSJR301 | r-pal | 0 | 2 | 13 | -40 | 9 | 6 | 10 |
| Model PsfiA-GFP threshold 6 | eSJR214 | l-pal | 0 | -1 | 25 | -53 | 5 | 2 | 8 |
| Model PsfiA-GFP threshold 6 | eSJR302 | 2-pal | 0 | 1 | 10 | -34 | 7 | 10 | 9 |
| Model PsfiA-GFP threshold 6 | eSJR206 | wt | 1 | -5 | 40 | -52 | 5 | 20 | 4 |
| Model PsfiA-GFP threshold 6 | eSJR206 | wt | 2 | -1 | 40 | -61 | 7 | 20 | 7 |
| Model PsfiA-GFP threshold 6 | eSJR206 | wt | 3 | 10 | 10 | -51 | 10 | 10 | 8 |

| | | | | alpha | err |
|---|---|---|---|---|---|
| Model PsfiA-GFP threshold 10 | eSJR206 | wt | 0 | 0,00053 | 0,00009 |
| Model PsfiA-GFP threshold 10 | eSJR301 | r-pal | 0 | 0,005 | 0,001 |
| Model PsfiA-GFP threshold 10 | eSJR214 | l-pal | 0 | 0,02 | 0,005 |
| Model PsfiA-GFP threshold 10 | eSJR302 | 2-pal | 0 | 0,026 | 0,005 |
| Model PsfiA-GFP threshold 10 | eSJR206 | wt | 1 | 0,0038 | 0,0009 |
| Model PsfiA-GFP threshold 10 | eSJR206 | wt | 2 | 0,012 | 0,003 |
| Model PsfiA-GFP threshold 10 | eSJR206 | wt | 3 | 0,021 | 0,004 |

| | | | | Relative distance to model [%] | | | error | | |
|---|---|---|---|---|---|---|---|---|---|
| | | | | gly | glu | glu+aa | gly | glu | glu+aa |
| Model PsfiA-GFP threshold 10 | eSJR206 | wt | 0 | 10 | 10 | -71 | 10 | 10 | 9 |
| Model PsfiA-GFP threshold 10 | eSJR301 | r-pal | 0 | 0 | 32 | -74 | 20 | 4 | 10 |
| Model PsfiA-GFP threshold 10 | eSJR214 | l-pal | 0 | -3 | 35 | -64 | 8 | 4 | 3 |
| Model PsfiA-GFP threshold 10 | eSJR302 | 2-pal | 0 | 2 | 30 | -55 | 7 | 10 | 9 |
| Model PsfiA-GFP threshold 10 | eSJR206 | wt | 1 | 0 | 40 | -65 | 5 | 10 | 7 |
| Model PsfiA-GFP threshold 10 | eSJR206 | wt | 2 | 5 | 30 | -77 | 10 | 20 | 5 |
| Model PsfiA-GFP threshold 10 | eSJR206 | wt | 3 | 10 | 0 | -60 | 10 | 10 | 10 |

| | | | | alpha | err |
|---|---|---|---|---|---|
| Model PsfiA-GFP threshold 12 | eSJR206 | wt | 0 | 0,0004 | 0,00007 |
| Model PsfiA-GFP threshold 12 | eSJR301 | r-pal | 0 | 0,0026 | 0,0009 |
| Model PsfiA-GFP threshold 12 | eSJR214 | l-pal | 0 | 0,016 | 0,004 |
| Model PsfiA-GFP threshold 12 | eSJR302 | 2-pal | 0 | 0,02 | 0,004 |
| Model PsfiA-GFP threshold 12 | eSJR206 | wt | 1 | 0,003 | 0,0007 |
| Model PsfiA-GFP threshold 12 | eSJR206 | wt | 2 | 0,009 | 0,002 |
| Model PsfiA-GFP threshold 12 | eSJR206 | wt | 3 | 0,016 | 0,003 |

| | | | | Relative distance to model [%] | | | error | | |
|---|---|---|---|---|---|---|---|---|---|
| | | | | gly | glu | glu+aa | gly | glu | glu+aa |
| Model PsfiA-GFP threshold 12 | eSJR206 | wt | 0 | 10 | 0 | -70 | 10 | 10 | 10 |
| Model PsfiA-GFP threshold 12 | eSJR301 | r-pal | 0 | 0 | 42 | -80 | 20 | 5 | 10 |
| Model PsfiA-GFP threshold 12 | eSJR214 | l-pal | 0 | -5 | 42 | -66 | 7 | 5 | 5 |
| Model PsfiA-GFP threshold 12 | eSJR302 | 2-pal | 0 | -1 | 30 | -57 | 9 | 10 | 9 |
| Model PsfiA-GFP threshold 12 | eSJR206 | wt | 1 | 2 | 40 | -73 | 6 | 10 | 5 |
| Model PsfiA-GFP threshold 12 | eSJR206 | wt | 2 | 10 | 30 | -81 | 10 | 20 | 3 |
| Model PsfiA-GFP threshold 12 | eSJR206 | wt | 3 | 10 | 0 | -60 | 10 | 10 | 20 |

| | | | | alpha | err |
|---|---|---|---|---|---|
| Model PsfiA-GFP threshold 14 | eSJR206 | wt | 0 | 0,0003 | 0,00006 |
| Model PsfiA-GFP threshold 14 | eSJR301 | r-pal | 0 | 0,0014 | 0,0005 |
| Model PsfiA-GFP threshold 14 | eSJR214 | l-pal | 0 | 0,012 | 0,003 |
| Model PsfiA-GFP threshold 14 | eSJR302 | 2-pal | 0 | 0,015 | 0,004 |
| Model PsfiA-GFP threshold 14 | eSJR206 | wt | 1 | 0,0023 | 0,0006 |
| Model PsfiA-GFP threshold 14 | eSJR206 | wt | 2 | 0,007 | 0,002 |
| Model PsfiA-GFP threshold 14 | eSJR206 | wt | 3 | 0,013 | 0,003 |

| | | | | Relative distance to model [%] | | | error | | |
|---|---|---|---|---|---|---|---|---|---|
| | | | | gly | glu | glu+aa | gly | glu | glu+aa |
| Model PsfiA-GFP threshold 14 | eSJR206 | wt | 0 | 10 | 0 | -74 | 10 | 10 | 8 |
| Model PsfiA-GFP threshold 14 | eSJR301 | r-pal | 0 | 0 | 44,2 | -84 | 20 | 0,7 | 8 |
| Model PsfiA-GFP threshold 14 | eSJR214 | l-pal | 0 | -8 | 53 | -67 | 8 | 6 | 5 |
| Model PsfiA-GFP threshold 14 | eSJR302 | 2-pal | 0 | -3 | 40 | -58 | 9 | 10 | 8 |
| Model PsfiA-GFP threshold 14 | eSJR206 | wt | 1 | -1 | 40 | -72 | 9 | 20 | 5 |
| Model PsfiA-GFP threshold 14 | eSJR206 | wt | 2 | 10 | 30 | -86 | 10 | 30 | 4 |
| Model PsfiA-GFP threshold 14 | eSJR206 | wt | 3 | 11 | 0 | -70 | 10 | 10 | 20 |
