## Supplementary data: Mother machine values for "Growth-dependent heterogeneity in the DNA damage response in *Escherichia coli*"

List of values as support in the main text

| Metric | strainID | Strain | media: gly | glu | glu+aa |
|---|---|---|---|---|---|
| Experiment ID | eSJR206 | WT | 20200201 | 20191020 | 20190818 |
| | eSJR206 | WT | 20200212 | 20200314 | 20190828 |
| | eSJR206 | WT | 20200221 | | 20200321 |
| | eSJR302 | 2pal | 20200201 | 20191020 | 20190818 |
| | eSJR302 | 2pal | 20200212 | 20191111 | 20190828 |
| | eSJR302 | 2pal | 20200221 | 20200314 | 20200321 |
| Imaging interval [h] | eSJR206 | WT | 0,13 | 0,13 | 0,083 |
| | eSJR206 | WT | 0,13 | 0,13 | 0,083 |
| | eSJR206 | WT | 0,2 | | 0,083 |
| | eSJR302 | 2pal | 0,13 | 0,13 | 0,083 |
| | eSJR302 | 2pal | 0,13 | 0,13 | 0,083 |
| | eSJR302 | 2pal | 0,2 | 0,13 | 0,083 |
| Tracking time [h] | eSJR206 | WT | 20 | 21 | 12 |
| | eSJR206 | WT | 11 | 17 | 13 |
| | eSJR206 | WT | 42 | | 15 |
| | eSJR302 | 2pal | 23 | 20 | 10 |
| | eSJR302 | 2pal | 22 | 22 | 15 |
| | eSJR302 | 2pal | 26 | 9,2 | 15 |
| Number of lineages | eSJR206 | WT | 45 | 56 | 83 |
| | eSJR206 | WT | 247 | 33 | 194 |
| | eSJR206 | WT | 212 | | 140 |
| | eSJR302 | 2pal | 92 | 366 | 172 |
| | eSJR302 | 2pal | 63 | 249 | 277 |
| | eSJR302 | 2pal | 448 | 284 | 160 |
| Number of division recorded | eSJR206 | WT | 163 | 506 | 1179 |
| | eSJR206 | WT | 611 | 341 | 3032 |
| | eSJR206 | WT | 2569 | | 2382 |
| | eSJR302 | 2pal | 151 | 1647 | 776 |
| | eSJR302 | 2pal | 106 | 2069 | 1824 |
| | eSJR302 | 2pal | 1115 | 517 | 1150 |

| Metric | strainID | Strain | avg | | | err | | |
|---|---|---|---|---|---|---|---|---|
| | | | gly | glu | glu+aa | gly | glu | glu+aa |
| Division time [h] | eSJR206 | WT | 3,7 | 1,56 | 0,63 | 1,5 | 0,97 | 0,3 |
| | eSJR206 | WT | 2,64 | 1,21 | 0,67 | 0,66 | 0,53 | 0,36 |
| | eSJR206 | WT | 2,8 | | 0,63 | 1,5 | | 0,32 |
| | eSJR302 | 2pal | 5,8 | 2,3 | 1,28 | 3,1 | 1,3 | 0,54 |
| | eSJR302 | 2pal | 5,5 | 1,5 | 1,37 | 2,4 | 0,81 | 0,59 |
| | eSJR302 | 2pal | 4,9 | 2,1 | 1,06 | 2,2 | 1 | 0,43 |
| Length at birth [µm] | eSJR206 | WT | 1,39 | 2,17 | 2,61 | 0,2 | 0,98 | 0,79 |
| | eSJR206 | WT | 1,7 | 1,67 | 2,47 | 0,21 | 0,25 | 0,68 |
| | eSJR206 | WT | 1,6 | | 2,33 | 0,22 | | 0,74 |
| | eSJR302 | 2pal | 1,58 | 2,8 | 2,1 | 0,24 | 2,3 | 1,3 |
| | eSJR302 | 2pal | 1,88 | 1,81 | 1,71 | 0,65 | 0,65 | 0,96 |
| | eSJR302 | 2pal | 1,88 | 2,6 | 2,6 | 0,47 | 1,8 | 1,9 |
| Length at division [µm] | eSJR206 | WT | 2,94 | 4,1 | 4,9 | 0,42 | 1,4 | 1,3 |
| | eSJR206 | WT | 3,39 | 3,3 | 4,7 | 0,4 | 1,2 | 1,3 |
| | eSJR206 | WT | 3,21 | | 4,4 | 0,41 | | 1,3 |
| | eSJR302 | 2pal | 3,38 | 5,5 | 4,1 | 0,49 | 3,6 | 2,4 |
| | eSJR302 | 2pal | 3,9 | 3,6 | 3,4 | 1 | 1,2 | 1,6 |
| | eSJR302 | 2pal | 3,81 | 4,7 | 5 | 0,69 | 2,7 | 2,8 |
| Elongation rate-length [/h] | eSJR206 | WT | 0,214 | 0,47 | 1,07 | 0,062 | 0,15 | 0,27 |
| | eSJR206 | WT | 0,255 | 0,57 | 1,04 | 0,05 | 0,14 | 0,28 |
| | eSJR206 | WT | 0,274 | | 1,09 | 0,084 | | 0,28 |
| | eSJR302 | 2pal | 0,144 | 0,321 | 0,58 | 0,055 | 0,093 | 0,19 |
| | eSJR302 | 2pal | 0,146 | 0,49 | 0,54 | 0,055 | 0,15 | 0,2 |
| | eSJR302 | 2pal | 0,156 | 0,314 | 0,66 | 0,052 | 0,086 | 0,13 |
| Elongation rate-length $R^2$ | eSJR206 | WT | 0,951 | 0,945 | 0,977 | 0,036 | 0,061 | 0,053 |
| | eSJR206 | WT | 0,963 | 0,951 | 0,977 | 0,026 | 0,069 | 0,042 |
| | eSJR206 | WT | 0,961 | | 0,976 | 0,042 | | 0,031 |
| | eSJR302 | 2pal | 0,93 | 0,949 | 0,958 | 0,12 | 0,077 | 0,052 |
| | eSJR302 | 2pal | 0,952 | 0,946 | 0,953 | 0,038 | 0,066 | 0,071 |
| | eSJR302 | 2pal | 0,955 | 0,93 | 0,972 | 0,055 | 0,093 | 0,04 |
| Cell-cycle mean PsfiA-GFP [a.u.] | eSJR206 | WT | 0,89 | 1,11 | 1,04 | 0,9 | 0,27 | 0,22 |
| | eSJR206 | WT | 0,84 | 0,88 | 0,96 | 0,44 | 0,47 | 0,24 |
| | eSJR206 | WT | 0,84 | | 1,01 | 0,41 | | 0,41 |
| | eSJR302 | 2pal | 0,9 | 1,34 | 1,28 | 0,22 | 0,34 | 0,43 |
| | eSJR302 | 2pal | 1,14 | 1,48 | 1,23 | 0,93 | 0,83 | 0,46 |
| | eSJR302 | 2pal | 0,97 | 1,14 | 1,44 | 0,39 | 0,5 | 0,49 |
| Cell-cycle mean PtetO-mKate [a.u.] | eSJR206 | WT | 1,7 | 1,44 | 0,73 | 0,29 | 0,29 | 0,28 |
| | eSJR206 | WT | 1,85 | 1,66 | 0,79 | 0,25 | 0,32 | 0,31 |
| | eSJR206 | WT | 1,98 | | 1,15 | 0,45 | | 0,48 |
| | eSJR302 | 2pal | 1,91 | 2,23 | 1,42 | 0,37 | 0,44 | 0,38 |
| | eSJR302 | 2pal | 2,19 | 1,95 | 1,48 | 0,39 | 0,45 | 0,35 |
| | eSJR302 | 2pal | 2,19 | 1,87 | 1,62 | 0,42 | 0,34 | 0,32 |

| | | | | | | avg | | | err | | |
|---|---|---|---|---|---|---|---|---|---|---|---|
| **Estimated population growth rate / Lambda1 [/h]** | eSJR206 | WT | 0,206138 | 0,53244 | 1,244227 | 0,256657 | 0,588624 | 1,222233 | 0,025318 | 0,056183 | 0,02526 |
| | eSJR206 | WT | 0,278917 | 0,644807 | 1,171849 | | | | | | |
| | eSJR206 | WT | 0,284915 | | 1,250625 | | | | | | |
| | eSJR302 | 2pal | 0,13496 | 0,351295 | 0,614416 | 0,143491 | 0,420874 | 0,634543 | 0,008531 | 0,056531 | 0,047145 |
| | eSJR302 | 2pal | 0,13496 | 0,53284 | 0,564831 | | | | | | |
| | eSJR302 | 2pal | 0,160552 | 0,378487 | 0,724383 | | | | | | |

| | | | | | | avg | | | err | | |
|---|---|---|---|---|---|---|---|---|---|---|---|
| **Estimated doubling-rate [dbl/h]** | eSJR206 | WT | 0,297395 | 0,768149 | 1,79504 | 0,370277 | 0,849204 | 1,76331 | 0,036527 | 0,081055 | 0,036443 |
| | eSJR206 | WT | 0,402392 | 0,93026 | 1,69062 | | | | | | |
| | eSJR206 | WT | 0,411045 | | 1,80427 | | | | | | |
| | eSJR302 | 2pal | 0,194706 | 0,506811 | 0,886415 | 0,207013 | 0,607193 | 0,915452 | 0,012307 | 0,081557 | 0,068016 |
| | eSJR302 | 2pal | 0,194706 | 0,768726 | 0,814879 | | | | | | |
| | eSJR302 | 2pal | 0,231628 | 0,546041 | 1,045064 | | | | | | |

| | | | | | |
|---|---|---|---|---|---|
| **Alpha SOS>5 [/h]** | eSJR302 | 2pal | 0,0091 | 0,017 | 0,033 |
| | eSJR302 | 2pal | 0,0116 | 0,028 | 0,029 |
| | eSJR302 | 2pal | 0,013 | 0,029 | 0,038 |

| | | | | | | avg | | | err | | |
|---|---|---|---|---|---|---|---|---|---|---|---|
| **Estimated f2 SOS>5 [fraction]** | eSJR302 | 2pal | 0,067427 | 0,048392 | 0,05371 | 0,078117 | 0,059187 | 0,052504 | 0,005535 | 0,008799 | 0,000684 |
| | eSJR302 | 2pal | 0,085952 | 0,052549 | 0,051343 | | | | | | |
| | eSJR302 | 2pal | 0,080971 | 0,076621 | 0,052458 | | | | | | |

**RAW DATA**

**Alpha estimates**

| strain | media | date | nbelow_SOS_5 | nabove_SOS_5 | alpha_maxlh_SOS_5 | alpha_p5_SOS_5 | alpha_p95_SOS_5 | nbelow_SOS_6 | nabove_SOS_6 | alpha_maxlh_SOS_6 | alpha_p5_SOS_6 | alpha_p95_SOS_6 | nbelow_SOS_7 | nabove_SOS_7 | alpha_maxlh_SOS_7 | alpha_p5_SOS_7 | alpha_p95_SOS_7 | nbelow_SOS_8 | nabove_SOS_8 | alpha_maxlh_SOS_8 | alpha_p5_SOS_8 | alpha_p95_SOS_8 | nbelow_SOS_9 | nabove_SOS_9 | alpha_maxlh_SOS_9 | alpha_p5_SOS_9 | alpha_p95_SOS_9 | nbelow_SOS_10 |
|---|---|---|---|---|---|---|---|---|---|---|---|---|---|---|---|---|---|---|---|---|---|---|---|---|---|---|---|---|
| wt | Gly | 20200201 | 43 | 1 | 0,00115 | 0,00051 | 0,00575 | 43 | 1 | 0,00115 | 0,0005 | 0,0053 | 43 | 1 | 0,00115 | 0,0005 | 0,00535 | 44 | 1 | 0,00112 | 0,0005 | 0,00538 | 44 | 1 | 0,00112 | 0,00049 | 0,00538 | 44 |
| wt | Gly | 20200212 | 237 | 8 | 0,0032 | 0,0019 | 0,0057 | 237 | 8 | 0,0032 | 0,0019 | 0,0057 | 238 | 7 | 0,0028 | 0,0016 | 0,0052 | 240 | 6 | 0,0024 | 0,0013 | 0,0047 | 240 | 6 | 0,0024 | 0,0013 | 0,0047 | 240 |
| wt | Gly | 20200221 | 195 | 15 | 0,0018 | 0,0012 | 0,0027 | 196 | 15 | 0,0018 | 0,0012 | 0,0027 | 199 | 13 | 0,00152 | 0,001 | 0,00241 | 201 | 11 | 0,00128 | 0,00083 | 0,00213 | 203 | 9 | 0,00105 | 0,00066 | 0,00184 | 203 |
| wt | Glu | 20191020 | 55 | 1 | 0,00097 | 0,00044 | 0,00465 | 55 | 1 | 0,00097 | 0,00044 | 0,00462 | 55 | 1 | 0,00097 | 0,00045 | 0,00482 | 55 | 1 | 0,00097 | 0,00043 | 0,00476 | 56 | 0 | 0 | 0,000075 | 0,003046 | 56 |
| wt | Glu | 20200314 | 31 | 2 | 0,0039 | 0,0016 | 0,0119 | 32 | 1 | 0,00194 | 0,00081 | 0,00925 | 32 | 1 | 0,00194 | 0,00078 | 0,00914 | 32 | 1 | 0,00194 | 0,00078 | 0,00881 | 32 | 1 | 0,00194 | 0,0008 | 0,00865 | 32 |
| wt | Gluaa | 20190818 | 83 | 0 | 0 | 0,000091 | 0,003466 | 83 | 0 | 0 | 0,000092 | 0,003392 | 83 | 0 | 0 | 0,000091 | 0,003416 | 83 | 0 | 0 | 0,000091 | 0,003495 | 83 | 0 | 0 | 0,000091 | 0,003475 | 83 |
| wt | Gluaa | 20190828 | 190 | 4 | 0,00158 | 0,00082 | 0,00362 | 192 | 2 | 0,00079 | 0,00037 | 0,0025 | 192 | 2 | 0,00079 | 0,00038 | 0,00253 | 192 | 2 | 0,00079 | 0,00037 | 0,00251 | 192 | 2 | 0,00079 | 0,00037 | 0,00251 | 192 |
| wt | Gluaa | 20200321 | 132 | 8 | 0,0044 | 0,0026 | 0,0079 | 133 | 7 | 0,0039 | 0,0022 | 0,0072 | 136 | 4 | 0,0022 | 0,0011 | 0,005 | 137 | 3 | 0,00162 | 0,0008 | 0,0042 | 138 | 2 | 0,00108 | 0,00051 | 0,00345 | 138 |
| 2pal | Gly | 20200201 | 70 | 15 | 0,0091 | 0,0061 | 0,0139 | 71 | 14 | 0,0084 | 0,0055 | 0,0131 | 72 | 14 | 0,0083 | 0,0054 | 0,013 | 73 | 13 | 0,0076 | 0,0049 | 0,012 | 74 | 12 | 0,007 | 0,0044 | 0,0113 | 76 |
| 2pal | Gly | 20200212 | 44 | 12 | 0,0116 | 0,0076 | 0,0191 | 44 | 12 | 0,0115 | 0,0073 | 0,0192 | 48 | 9 | 0,0083 | 0,0051 | 0,0146 | 49 | 8 | 0,0074 | 0,0043 | 0,0129 | 50 | 7 | 0,0064 | 0,0036 | 0,012 | 51 |
| 2pal | Gly | 20200221 | 310 | 123 | 0,013 | 0,011 | 0,015 | 320 | 113 | 0,012 | 0,01 | 0,014 | 327 | 106 | 0,0112 | 0,0096 | 0,0131 | 336 | 97 | 0,0102 | 0,0086 | 0,012 | 341 | 92 | 0,0096 | 0,0081 | 0,0114 | 351 |
| 2pal | Glu | 20191020 | 262 | 94 | 0,017 | 0,014 | 0,02 | 267 | 90 | 0,016 | 0,013 | 0,019 | 277 | 80 | 0,014 | 0,011 | 0,016 | 280 | 77 | 0,013 | 0,011 | 0,016 | 294 | 65 | 0,0108 | 0,0088 | 0,0133 | 303 |
| 2pal | Glu | 20191111 | 134 | 113 | 0,028 | 0,024 | 0,033 | 139 | 108 | 0,026 | 0,022 | 0,031 | 148 | 99 | 0,024 | 0,02 | 0,028 | 157 | 90 | 0,021 | 0,018 | 0,025 | 164 | 83 | 0,019 | 0,016 | 0,023 | 170 |
| 2pal | Glu | 20200314 | 208 | 60 | 0,029 | 0,024 | 0,036 | 216 | 52 | 0,025 | 0,02 | 0,031 | 218 | 50 | 0,024 | 0,019 | 0,03 | 224 | 45 | 0,021 | 0,017 | 0,027 | 229 | 41 | 0,019 | 0,015 | 0,025 | 238 |
| 2pal | Gluaa | 20190818 | 118 | 45 | 0,033 | 0,026 | 0,041 | 124 | 40 | 0,029 | 0,023 | 0,037 | 128 | 37 | 0,026 | 0,02 | 0,035 | 129 | 37 | 0,026 | 0,019 | 0,034 | 132 | 34 | 0,023 | 0,018 | 0,031 | 135 |
| 2pal | Gluaa | 20190828 | 178 | 96 | 0,029 | 0,025 | 0,035 | 187 | 88 | 0,027 | 0,022 | 0,032 | 194 | 81 | 0,024 | 0,02 | 0,029 | 201 | 74 | 0,022 | 0,018 | 0,026 | 205 | 71 | 0,021 | 0,017 | 0,025 | 209 |
| 2pal | Gluaa | 20200321 | 93 | 62 | 0,038 | 0,031 | 0,047 | 98 | 57 | 0,035 | 0,028 | 0,044 | 102 | 53 | 0,032 | 0,026 | 0,04 | 105 | 50 | 0,029 | 0,023 | 0,038 | 106 | 50 | 0,029 | 0,023 | 0,037 | 110 |

| nabove_SOS_10 | alpha_maxlh_SOS_10 | alpha_p5_SOS_10 | alpha_p95_SOS_10 | nbelow_SOS_11 | nabove_SOS_11 | alpha_maxlh_SOS_11 | alpha_p5_SOS_11 | alpha_p95_SOS_11 | nbelow_SOS_12 | nabove_SOS_12 | alpha_maxlh_SOS_12 | alpha_p5_SOS_12 | alpha_p95_SOS_12 | nbelow_SOS_13 | nabove_SOS_13 | alpha_maxlh_SOS_13 | alpha_p5_SOS_13 | alpha_p95_SOS_13 | nbelow_SOS_14 | nabove_SOS_14 | alpha_maxlh_SOS_14 | alpha_p5_SOS_14 | alpha_p95_SOS_14 | nbelow_SOS_15 | nabove_SOS_15 | alpha_maxlh_SOS_15 | alpha_p5_SOS_15 | alpha_p95_SOS_15 | nbelow_SOS_16 | nabove_SOS_16 | alpha_maxlh_SOS_16 | alpha_p5_SOS_16 | alpha_p95_SOS_16 |
|---|---|---|---|---|---|---|---|---|---|---|---|---|---|---|---|---|---|---|---|---|---|---|---|---|---|---|---|---|---|---|---|---|---|
| 1 | 0,00112 | 0,0005 | 0,00536 | 44 | 1 | 0,00112 | 0,0005 | 0,00538 | 44 | 1 | 0,00112 | 0,0005 | 0,00533 | 44 | 1 | 0,00112 | 0,00048 | 0,00522 | 45 | 0 | 0 | 0,000091 | 0,003454 | 45 | 0 | 0 | 0,000091 | 0,003568 | 45 | 0 | 0 | 0,000095 | 0,003507 |
| 6 | 0,0024 | 0,0013 | 0,0046 | 240 | 6 | 0,0023 | 0,0013 | 0,0047 | 240 | 7 | 0,0027 | 0,0016 | 0,0052 | 242 | 5 | 0,0019 | 0,001 | 0,0041 | 242 | 5 | 0,0019 | 0,001 | 0,004 | 242 | 5 | 0,0019 | 0,001 | 0,0041 | 243 | 4 | 0,00155 | 0,0008 | 0,00351 |
| 9 | 0,00104 | 0,00065 | 0,00183 | 204 | 8 | 0,00093 | 0,00057 | 0,00167 | 204 | 8 | 0,00093 | 0,00057 | 0,00169 | 205 | 7 | 0,00081 | 0,00049 | 0,00154 | 206 | 6 | 0,00069 | 0,00041 | 0,0014 | 206 | 6 | 0,00069 | 0,00041 | 0,00139 | 206 | 6 | 0,00069 | 0,00041 | 0,00139 |
| 0 | 0 | 0,000081 | 0,003146 | 56 | 0 | 0 | 0,000081 | 0,003193 | 56 | 0 | 0 | 0,00008 | 0,003081 | 56 | 0 | 0 | 0,000079 | 0,003076 | 56 | 0 | 0 | 0,000077 | 0,003235 | 56 | 0 | 0 | 0,00008 | 0,003227 | 56 | 0 | 0 | 0,000079 | 0,003069 |
| 1 | 0,00194 | 0,00081 | 0,0097 | 32 | 1 | 0,00194 | 0,00078 | 0,00925 | 32 | 1 | 0,00194 | 0,00079 | 0,00882 | 32 | 1 | 0,00194 | 0,00079 | 0,00928 | 32 | 1 | 0,00194 | 0,00082 | 0,009 | 32 | 1 | 0,00194 | 0,0008 | 0,0097 | 33 | 0 | 0 | 0,00016 | 0,00594 |
| 0 | 0 | 0,000091 | 0,003485 | 83 | 0 | 0 | 0,000089 | 0,003321 | 83 | 0 | 0 | 0,00009 | 0,00339 | 83 | 0 | 0 | 0,000092 | 0,003449 | 83 | 0 | 0 | 0,00009 | 0,003463 | 83 | 0 | 0 | 0,000091 | 0,003434 | 83 | 0 | 0 | 0,000091 | 0,003526 |
| 2 | 0,00079 | 0,00038 | 0,00256 | 192 | 2 | 0,00078 | 0,00037 | 0,0025 | 192 | 2 | 0,00078 | 0,00037 | 0,00249 | 192 | 2 | 0,00078 | 0,00037 | 0,00252 | 192 | 2 | 0,00078 | 0,00037 | 0,00252 | 194 | 0 | 0 | 0,000027 | 0,001306 | 194 | 0 | 0 | 0,000028 | 0,001327 |
| 2 | 0,00108 | 0,00051 | 0,00347 | 139 | 1 | 0,00054 | 0,00024 | 0,00264 | 139 | 1 | 0,00054 | 0,00025 | 0,00271 | 140 | 0 | 0 | 0,000038 | 0,00178 | 140 | 0 | 0 | 0,00004 | 0,001788 | 140 | 0 | 0 | 0,000039 | 0,001763 | 140 | 0 | 0 | 0,00004 | 0,001798 |
| 10 | 0,0057 | 0,0035 | 0,0096 | 79 | 7 | 0,004 | 0,0022 | 0,0074 | 80 | 6 | 0,0034 | 0,0019 | 0,0066 | 80 | 6 | 0,0034 | 0,0019 | 0,0066 | 80 | 6 | 0,0034 | 0,0019 | 0,0067 | 81 | 5 | 0,0028 | 0,0015 | 0,006 | 82 | 4 | 0,0023 | 0,0011 | 0,0051 |
| 6 | 0,0055 | 0,003 | 0,0111 | 51 | 6 | 0,0055 | 0,003 | 0,0106 | 52 | 5 | 0,0045 | 0,0023 | 0,0091 | 52 | 5 | 0,0045 | 0,0024 | 0,0092 | 53 | 4 | 0,0036 | 0,0018 | 0,0083 | 53 | 4 | 0,0036 | 0,0017 | 0,0081 | 53 | 5 | 0,0044 | 0,0023 | 0,0094 |
| 82 | 0,0084 | 0,0071 | 0,0102 | 356 | 77 | 0,0079 | 0,0066 | 0,0095 | 361 | 72 | 0,0073 | 0,006 | 0,0089 | 367 | 66 | 0,0066 | 0,0055 | 0,0082 | 373 | 60 | 0,006 | 0,0049 | 0,0074 | 379 | 54 | 0,0054 | 0,0043 | 0,0067 | 384 | 49 | 0,0049 | 0,0039 | 0,0062 |
| 56 | 0,0091 | 0,0074 | 0,0114 | 309 | 50 | 0,0081 | 0,0064 | 0,0102 | 315 | 44 | 0,007 | 0,0055 | 0,0091 | 322 | 37 | 0,0058 | 0,0045 | 0,0077 | 328 | 31 | 0,0049 | 0,0036 | 0,0066 | 336 | 23 | 0,0036 | 0,0026 | 0,0051 | 337 | 22 | 0,0034 | 0,0024 | 0,0049 |
| 78 | 0,018 | 0,015 | 0,021 | 179 | 69 | 0,015 | 0,013 | 0,019 | 187 | 61 | 0,013 | 0,011 | 0,017 | 194 | 54 | 0,0116 | 0,0093 | 0,0146 | 198 | 50 | 0,0106 | 0,0085 | 0,0135 | 202 | 46 | 0,0097 | 0,0077 | 0,0125 | 206 | 42 | 0,0088 | 0,0068 | 0,0114 |
| 32 | 0,014 | 0,011 | 0,019 | 239 | 31 | 0,014 | 0,01 | 0,019 | 244 | 28 | 0,0124 | 0,0092 | 0,0171 | 249 | 23 | 0,0101 | 0,0073 | 0,0142 | 252 | 20 | 0,0087 | 0,0062 | 0,0126 | 255 | 17 | 0,0073 | 0,005 | 0,0111 | 255 | 18 | 0,0078 | 0,0053 | 0,0116 |
| 31 | 0,021 | 0,016 | 0,028 | 137 | 30 | 0,02 | 0,015 | 0,027 | 142 | 25 | 0,016 | 0,012 | 0,023 | 146 | 21 | 0,0135 | 0,0097 | 0,0193 | 148 | 19 | 0,0122 | 0,0086 | 0,0179 | 150 | 18 | 0,0114 | 0,0079 | 0,017 | 151 | 17 | 0,0107 | 0,0075 | 0,0162 |
| 67 | 0,019 | 0,016 | 0,024 | 216 | 60 | 0,017 | 0,014 | 0,021 | 220 | 56 | 0,016 | 0,013 | 0,02 | 225 | 51 | 0,014 | 0,011 | 0,018 | 226 | 50 | 0,014 | 0,011 | 0,017 | 229 | 47 | 0,013 | 0,01 | 0,017 | 232 | 44 | 0,012 | 0,0093 | 0,0154 |
| 46 | 0,026 | 0,021 | 0,034 | 111 | 45 | 0,026 | 0,02 | 0,032 | 115 | 41 | 0,023 | 0,018 | 0,029 | 118 | 38 | 0,021 | 0,016 | 0,027 | 119 | 38 | 0,021 | 0,016 | 0,027 | 120 | 38 | 0,021 | 0,016 | 0,027 | 124 | 34 | 0,018 | 0,014 | 0,024 |

**Mother machine data**

| strain | media | date | interval | tracking_time | n_lineages | n_lineages_last_div_rmv | n_lineages_truncated_odd_growth | n_divisions | division_time_avg | division_time_std | length_at_birth_avg | length_at_birth_std | length_at_division_avg | length_at_division_std | length_grrate_avg | length_grrate_std | length_grrate_r2_avg | length_grrate_r2_std | cellcycle_mean_gfp_avg | cellcycle_mean_gfp_std | cellcycle_mean_mkate_avg | cellcycle_mean_mkate_std | estimated_pop_grates | average lambda 1 | average doubling rate |
|---|---|---|---|---|---|---|---|---|---|---|---|---|---|---|---|---|---|---|---|---|---|---|---|---|---|
| wt | Gly | 20200201 | 0,13 | 20 | 45 | 3 | 3 | 163 | 3,7 | 1,5 | 1,39 | 0,2 | 2,94 | 0,42 | 0,214 | 0,062 | 0,951 | 0,036 | 0,89 | 0,9 | 1,7 | 0,29 | 0,206138 | 0,256657 | 0,370277 |
| wt | Gly | 20200212 | 0,13 | 11 | 247 | 179 | 34 | 611 | 2,64 | 0,66 | 1,7 | 0,21 | 3,39 | 0,4 | 0,255 | 0,05 | 0,963 | 0,026 | 0,84 | 0,44 | 1,85 | 0,25 | 0,278917 | | |
| wt | Gly | 20200221 | 0,2 | 42 | 212 | 45 | 40 | 2569 | 2,8 | 1,5 | 1,6 | 0,22 | 3,21 | 0,41 | 0,274 | 0,084 | 0,961 | 0,042 | 0,84 | 0,41 | 1,98 | 0,45 | 0,284915 | | |
| wt | Glu | 20191020 | 0,13 | 21 | 56 | 31 | 14 | 506 | 1,56 | 0,97 | 2,17 | 0,98 | 4,1 | 1,4 | 0,47 | 0,15 | 0,945 | 0,061 | 1,11 | 0,27 | 1,44 | 0,29 | 0,53244 | 0,588624 | 0,849204 |
| wt | Glu | 20200314 | 0,13 | 17 | 33 | 10 | 6 | 341 | 1,21 | 0,53 | 1,67 | 0,25 | 3,3 | 1,2 | 0,57 | 0,14 | 0,951 | 0,069 | 0,88 | 0,47 | 1,66 | 0,32 | 0,644807 | | |
| wt | Gluaa | 20190818 | 0,083 | 12 | 83 | 41 | 21 | 1179 | 0,63 | 0,3 | 2,61 | 0,79 | 4,9 | 1,3 | 1,07 | 0,27 | 0,977 | 0,053 | 1,04 | 0,22 | 0,73 | 0,28 | 1,244227 | 1,222233 | 1,76331 |
| wt | Gluaa | 20190828 | 0,083 | 13 | 194 | 66 | 29 | 3032 | 0,67 | 0,36 | 2,47 | 0,68 | 4,7 | 1,3 | 1,04 | 0,28 | 0,977 | 0,042 | 0,96 | 0,24 | 0,79 | 0,31 | 1,171849 | | |
| wt | Gluaa | 20200321 | 0,083 | 15 | 140 | 72 | 36 | 2382 | 0,63 | 0,32 | 2,33 | 0,74 | 4,4 | 1,3 | 1,09 | 0,28 | 0,976 | 0,031 | 1,01 | 0,41 | 1,15 | 0,48 | 1,250625 | | |
| 2pal | Gly | 20200201 | 0,13 | 23 | 92 | 7 | 18 | 151 | 5,8 | 3,1 | 1,58 | 0,24 | 3,38 | 0,49 | 0,144 | 0,055 | 0,93 | 0,12 | 0,9 | 0,22 | 1,91 | 0,37 | 0,13496 | 0,143491 | 0,207013 |
| 2pal | Gly | 20200212 | 0,13 | 22 | 63 | 4 | 15 | 106 | 5,5 | 2,4 | 1,88 | 0,65 | 3,9 | 1 | 0,146 | 0,055 | 0,952 | 0,038 | 1,14 | 0,93 | 2,19 | 0,39 | 0,13496 | | |
| 2pal | Gly | 20200221 | 0,2 | 26 | 448 | 30 | 92 | 1115 | 4,9 | 2,2 | 1,88 | 0,47 | 3,81 | 0,69 | 0,156 | 0,052 | 0,955 | 0,055 | 0,97 | 0,39 | 2,19 | 0,42 | 0,160552 | | |
| 2pal | Glu | 20191020 | 0,13 | 20 | 366 | 108 | 75 | 1647 | 2,3 | 1,3 | 2,8 | 2,3 | 5,5 | 3,6 | 0,321 | 0,093 | 0,949 | 0,077 | 1,34 | 0,34 | 2,23 | 0,44 | 0,351295 | 0,420874 | 0,607193 |
| 2pal | Glu | 20191111 | 0,13 | 22 | 249 | 59 | 61 | 2069 | 1,5 | 0,81 | 1,81 | 0,65 | 3,6 | 1,2 | 0,49 | 0,15 | 0,946 | 0,066 | 1,48 | 0,83 | 1,95 | 0,45 | 0,53284 | | |
| 2pal | Glu | 20200314 | 0,13 | 9,2 | 284 | 34 | 54 | 517 | 2,1 | 1 | 2,6 | 1,8 | 4,7 | 2,7 | 0,314 | 0,086 | 0,93 | 0,093 | 1,14 | 0,5 | 1,87 | 0,34 | 0,378487 | | |
| 2pal | Gluaa | 20190818 | 0,083 | 10 | 172 | 20 | 31 | 776 | 1,28 | 0,54 | 2,1 | 1,3 | 4,1 | 2,4 | 0,58 | 0,19 | 0,958 | 0,052 | 1,28 | 0,43 | 1,42 | 0,38 | 0,614416 | 0,634543 | 0,915452 |
| 2pal | Gluaa | 20190828 | 0,083 | 15 | 277 | 51 | 33 | 1824 | 1,37 | 0,59 | 1,71 | 0,96 | 3,4 | 1,6 | 0,54 | 0,2 | 0,953 | 0,071 | 1,23 | 0,46 | 1,48 | 0,35 | 0,564831 | | |
| 2pal | Gluaa | 20200321 | 0,083 | 15 | 160 | 40 | 41 | 1150 | 1,06 | 0,43 | 2,6 | 1,9 | 5 | 2,8 | 0,66 | 0,13 | 0,972 | 0,04 | 1,44 | 0,49 | 1,62 | 0,32 | 0,724383 | | |
